# The collective effect of non-nicotine constituents on the reinforcing and neuropharmacological effects of comprehensive cigarette smoke and electronic nicotine delivery system (ENDS) aerosol extracts in rats

**DOI:** 10.64898/2026.11.01.763124

**Authors:** MG LeSage, P Muelken, S Howard, S Wilde, A Saykao, JR Smethells, A Alcheva, T Olson, FM Leslie, S Lotfipour, KJ Kellar, I Stepanov, AC Harris

## Abstract

Tobacco product extracts containing nicotine and non-nicotine constituents are useful for understanding the collective role of these constituents in tobacco use disorder (TUD). However, few studies have used extracts with both water-soluble and non-water-soluble constituents with accompanying chemical analysis, or compared behavioral and neuropharmacological effects across product types. To address these limitations, we compared the relative abuse liability of cigarette smoke (CS) and electronic nicotine delivery system (ENDS) aerosol (EA) extracts to nicotine alone using intravenous self-administration in rats, and examined effects on nucleus accumbens (NAc) dopamine (DA) release and binding/function at nicotinic acetylcholine receptors (nAChRs) and other addiction-related receptors in vitro. CS extract contained higher levels of several behaviorally active non-nicotine constituents than EA extract. Both extracts produced a downward/leftward shift in the descending limb of the unit dose-response curve relative to nicotine alone, while only CS extract produced greater compensatory increases in responding compared to nicotine alone. Demand intensity (Q0) was lower for CS but not EA extract, relative to nicotine alone, whereas demand elasticity for both extracts was similar to nicotine alone. Stress-induced reinstatement was comparable between CS extract and nicotine alone but attenuated for EA extract. CS extract, but not EA extract, showed greater potency for α4β2 nAChR activation and substantial binding across numerous non-nAChRs (DA, serotonergic, opioid) relative to nicotine alone at a high screening concentration. CS extract, but not nicotine alone, also inhibited cyclooxygenase (COX)-1 and -2 at smoking-relevant concentrations. There was no difference in NAc DA release between formulations. These findings indicate non-nicotine constituents contribute to certain reinforcing and neuropharmacological effects of CS, and to a lesser extent EA. Our finding of COX inhibition by CS extract warrants further investigation as a potential mechanism contributing to the effects of CS and TUD.

## 1. Introduction

Tobacco products with greater abuse liability generally have higher nicotine content, produce higher nicotine serum levels, and are associated with faster nicotine absorption (Benowitz, 2008; Fant et al., 1999; Hatsukami et al., 2007; Houezec, 2003; Stiles et al., 2017). However, several findings indicate that factors other than nicotine contribute to product abuse liability. First, nicotine *<u>alone</u>*has limited abuse potential compared to other drugs in humans and animals (Foll and Goldberg, 2009; Manzardo et al., 2002; Rose et al., 2000). Second, pharmacotherapies that principally target nicotine’s mechanisms of action (e.g., varenicline, bupropion) have limited efficacy for smoking and electronic nicotine delivery system (ENDS) use cessation, suggesting they may fail to address other important mechanisms mediating smoking and ENDS use (Etter and Stapleton, 2006; Gómez-Coronado et al., 2018; Sahr et al., 2021; Stead et al., 2012). Third, smoke from very-low-nicotine-content (VLN, “denicotinized”) cigarettes is reinforcing in smokers, can be preferred over i.v. infusion of nicotine alone, is a better substitute for conventional cigarettes than NRT, increases D2 dopamine and µ-opioid receptor activation in numerous addiction-related brain regions, and can alleviate craving and anxiety during withdrawal as much as conventional cigarettes (Brody et al., 2008, 2009; Domino and Hirasawa-Fujita, 2018; Donny et al., 2007; Johnson et al., 2004; Nuechterlein et al., 2016; Pickworth et al., 1999; Rose et al., 2010b; Scott et al., 2007; Shahan et al., 2001). In addition, switching smokers to VLN cigarettes with a nicotine yield ≈ 95-98% lower than conventional cigarettes results in modest reductions in cigarettes smoked per day (CPD) compared to baseline usual-brand CPD if alternative nicotine products are not made freely available (Benowitz et al., 2012; Donny et al., 2015, 2007). Differences in nicotine content and delivery may therefore not fully account for differences in abuse liability between cigarettes and NCTPs.

Although these findings could be attributable to several environmental / sensory factors (e.g., conditioned reinforcing effects of nicotine-paired stimuli), several studies have shown that certain non-nicotine constituents in cigarette smoke and ENDS aerosol (e.g., monoamine oxidase (MAO) inhibitors, minor tobacco alkaloids, volatile organic compounds (VOCs) such as acetaldehyde, toluene, and benzene) have addiction-related CNS and behavioral effects (Bardo et al., 1999; Belluzzi et al., 2005; Bodnar et al., 2012; Caine et al., 2014; Goniewicz et al., 2014; Harris et al., 2025, 2019, 2015; Pazo et al., 2016; Polzin et al., 2007; Smethells et al., 2026, 2025; Smith et al., 2015; Stepanov et al., 2008; Truman et al., 2024). These constituents may therefore collectively contribute to the abuse liability of tobacco products and facilitate tobacco use disorder (TUD) by either enhancing the addiction-related effects of nicotine, producing addiction-related effects of their own, or both. As such, the limited success of current medications and VLN cigarettes to reduce TUD may be due, in part, to their failure to address the effects of non-nicotine constituents in tobacco products. The purpose of the present study was to advance further our understanding of the collective role of non-nicotine constituents in the abuse liability of tobacco products and examine potential mechanisms mediating their effects.

The present study addresses several limitations of prior preclinical research on the TUD-related effects of non-nicotine constituents. First, most animal studies have only evaluated nicotine and/or other known constituents in isolation. This approach alone is insufficient to examine mechanisms of TUD because as yet unidentified compounds may contribute positively or negatively to TUD, and it is the net effect of all compounds in a product that determines its actual abuse liability. One approach to address this issue has been examining the behavioral and CNS effects of inhalational exposure to cigarette smoke (CS) or ENDS aerosol (EA, (Brennan et al., 2014; Cooper et al., 2020, 2023; Foll et al., 2022; Gellner et al., 2016b; LeSage et al., 2018). Although this approach mimics the route used in humans and its peripheral sensory effects (taste, smell), its utility for discerning CNS mechanisms of TUD is limited because a) it does not allow isolation of the CNS effects of CS or EA from its peripheral sensory effects, b) the considerable stress of CS exposure can have widespread CNS effects that may be unrelated to TUD (Brennan et al., 2014), c) accurate dosing can be difficult, and d) some key models (e.g., CS self-administration) are not feasible. Another approach that addresses these limitations of inhalational exposure has been parenteral administration of extracts prepared from tobacco, ENDS liquids, CS, or EA (Brennan et al., 2014; Gellner et al., 2016b; LeSage et al., 2018; Marusich and Palmatier, 2023). Several studies from our lab and others evaluating effects of CS and EA extracts have reported greater addiction-related behavioral and neurobiological effects of these extracts than nicotine alone (Ambrose et al., 2007; Brennan et al., 2014; Costello et al., 2014; Danielson et al., 2014; Harris et al., 2022, 2020; Lee et al., 2016; LeSage et al., 2018; Touiki et al., 2007), suggesting non-nicotine constituents contribute to the addiction-related effects of the extracts. As such, the study of CS and EA extracts in animals is a vital translational model of tobacco product use allowing comparison of the abuse liability of different classes of tobacco products and discovery of their unique TUD-related mechanisms of action.

A second limitation of prior studies is that they have typically used extracts of water-soluble constituents in the particulate phase of CS and EA, excluding potentially important constituents that are not water soluble and/or occur in the gas phase of CS or EA (e.g., abused VOCs like toluene). Moreover, few, if any, constituents in CS and EA extracts have been measured in behavioral studies (Brennan et al., 2015; K A Brennan et al., 2013; Katharine A Brennan et al., 2013; Lee et al., 2016; Marusich and Palmatier, 2023), preventing identification of the non-nicotine constituents contributing to the behavioral effects of CS and EA and preventing understanding of similarities/differences in extract effects across studies(Costello et al., 2014; Gellner et al., 2016a). It also precludes detection of novel harmful and potentially harmful constituents (HPHCs). The FDA’s list of HPHCs is critical to tobacco regulation because it is used to a) identify constituents that may require product standards, b) determine the potential risk of new products, and c) inform the public of the risks of those and other products (Donny et al., 2012; LeSage et al., 2018).

A third limitation of prior preclinical research is that studies of the CNS mechanisms mediating the abuse liability of tobacco product extracts have been limited. Regarding CS extracts, studies show they have greater effects than nicotine alone on several TUD-related neurobiological processes, including nAChR upregulation (Ambrose et al., 2007; Cano et al., 2020), dopamine (DA) reuptake (Carr et al., 1992; Danielson et al., 2014), serotonin (5-HT) neuron firing (Touiki et al., 2007), and striatal glutamate release (Ryu et al., 2018). In addition, self-administration (SA) of CS extracts can be more resistant to reduction by nAChR and 5-HT antagonists than SA of nicotine alone (Katharine A Brennan et al., 2013; Costello et al., 2014). Effects of CS extract on other neural processes (e.g. MAO inhibition) have been mixed (Carr and Basham, 1991; Costello et al., 2014; Harris et al., 2022; G. Liu et al., 2022; Marti et al., 2011). The neurobiological effects of EA extracts have received little attention, however we have shown no differences in nAChR binding or functional activity between nicotine-containing e-liquids and nicotine alone (Harris et al., 2018; M G LeSage et al., 2016), while others have shown certain e-liquids can inhibit MAO A and B in vitro (Truman et al., 2019)

To address these limitations, we have developed comprehensive CS and EA extracts that include water-soluble and non-water-soluble constituents from both the particulate and gas phase of CS and EA from two popular conventional products, Marlboro Gold cigarettes and Vuse ENDS. We have also measured a range of behaviorally relevant non-nicotine constituents in these extracts, such as minor alkaloids (e.g., nornicotine, anabasine), MAO inhibitors (e.g., the β-carbolines harmane and norharmane), carbonyl VOCs (e.g. acetaldehyde, acetone), and other VOCs (e.g., benzene, toluene). Using these extracts, we previously found that acute injection of CS extract, but not EA extract, produced greater aversive / anhedonic effects in an intracranial self-stimulation model compared to nicotine alone at a high nicotine dose (1.0 mg/kg). We also found that CS extract but not nicotine alone or EA extract inhibited MAO in vitro, whereas neither formulation inhibited MAO ex vivo (Harris et al., 2022). These findings are consistent with another CS extract study (Costello et al., 2014) and suggest that MAO inhibition produced by CS extract was reversible, such that inhibition was detectable *in vitro* but likely lost during brain homogenization for ex vivo analysis.

A goal of the current study was to further characterize the TUD-related behavioral effects of CS and EA extracts by examining their reinforcing effects in an intravenous SA model in rats. This model is often considered the “gold standard” model of abuse liability in animals because it involves voluntary drug taking as occurs in humans. We also examined effects of nicotine alone and extracts on nucleus accumbens (NAc) dopamine (DA) release nAChR binding and function in vitro, critical neurobiological mechanisms associated with nicotine’s reinforcing effects. Importantly, given the complexity of constituents in extracts, with some having mechanisms of action that differ from nicotine, we compared extracts and nicotine alone in terms of binding and function at a range of addiction-related non-nAChRs (e.g. DA, 5-HT, GABA etc.) and enzymatic activity (e.g. acetylcholinesterase, cyclooxygenase, phosphodiesterase, etc.). Finally, given that sex differences in nicotine’s TUD-related behavioral effects have been reported in some studies (Chen et al., 2007; Lynch, 2009; Matos-Ocasio et al., 2021; Sanchez et al., 2013; Torres et al., 2009; Xue et al., 2018) and very few studies have examined sex differences in the effects of tobacco product extracts or non-nicotine constituents, both sexes were included in the present study to provide a preliminary evaluation of the impact of this variable.

## 2. Method

### 2.1. Animals

For self-administration studies, experimentally-naïve male and female adult Sprague-Dawley rats (Inotiv (formerly Envigo), Indianapolis, IN) aged 60-64 days at arrival were used. All rats were individually housed in a temperature-and humidity-controlled colony room with unlimited access to food and water under a reversed 12-h light/dark cycle (lights off at 10:00 hr) for one week prior to surgery and during a three- to six-day postoperative recovery period. Rats were then food restricted (16-18 or 18-20 grams/day for females and males, respectively) for the remainder of the protocol to facilitate operant performance and avoid detrimental health effects of long-term ad libitum feeding. Protocols were approved by the Institutional Animal Care and Use Committee of the Hennepin Healthcare Research Institute and were in accordance with the National Research Council’s Guide for the Care and Use of Laboratory Animals.

For microdialysis studies, male and female Sprague-Dawley rats were obtained from Charles River at postnatal (P) P80. After surgery, all animals were single-housed and maintained on a 12-h light/dark cycle (lights on at 07:00 am) with food and water available ad libitum. Rats were allowed at least 3 days of postoperative recovery before any treatments began. All tests were performed during the light part of the light-dark cycle. The study was conducted according to the guidelines of the Institutional Animal Care and Use Committee (IACUC) at the University of California, Irvine (UCI)

### 2.2. Apparatus

Standard operant conditioning chambers for rats (model ENV-007, Med Associates, Inc (St. Albans, VT) were used. Two response levers were located on the front wall 6 cm above the chamber floor on either side of an aperture for food delivery (not used in this study) located 2 cm above the floor. LED stimulus lights were located 2 cm above each response lever. Each chamber was placed inside a sound-attenuating cubicle equipped with an exhaust fan that provided masking noise. Infusion pumps (model PHM-100, Med Associates) placed outside each cubicle delivered infusions through Tygon tubing connected to a fluid swivel mounted above the chamber, and from the swivel through a spring leash connected to a vascular access harness or button (VAH95AB or VABR1B/22, respectively, Instech Laboratories, Plymouth Meeting, PA) mounted on the back of the rat. MED-PC IV or V software (Med Associates) was used for operating the apparatus and recording data.

### 2.3. Drugs

Nicotine-alone solutions consisted of (-)-nicotine (Sigma Chemical Co., St. Louis, MO) dissolved in sterile saline. CS and EA extracts were prepared as we have described previously (Harris et al. 2022). Briefly, CS extracts were prepared from Marlboro Gold cigarettes purchased in the Minneapolis area. Cigarettes were smoked on a Borgwaldt LX1 linear single port smoking machine under the Canadian Intense smoking regimen (55-ml puff volume, 2-puff duration, 30-s puff interval, and 100% blockade of filter ventilation holes). Total particulate matter (TPM) was collected on Cambridge filter pads, while gas phase constituents were trapped in an impinger containing 10 ml 50:50 saline/ETOH maintained at −23° C. After a total of 40 cigarettes were smoked (five cigarettes per filter pad), filter pads were extracted in 10 ml of a 50:50 saline/ETOH mixture. Use of ETOH for extraction captures a range of constituents, including those that are volatile and/or non-water soluble. ETOH has also been used as a solvent in preclinical studies from our lab and others evaluating addiction-related effects of extracts (Brennan et al., 2015; K A Brennan et al., 2013; Katharine A Brennan et al., 2013; Harris et al., 2022). The TPM fraction was cooled to -20° C and combined with the gas phase constituents to make the final CS extract (volume ≈ 17.5 ml). Levels of select constituents in undiluted extract are presented in Table 1.

**Table 1.** Levels of measured constituents in CS and EA extracts. Each value is the mean (±SEM) of 8-12 extract batches, except for other VOCs and solvents, which are from a single batch (measured in duplicate) and 5 batches, respectively.

|  | **CS Extract** | **EA Extract** |
|---|---|---|
| Nicotine (mg/ml) | 3.21 (±0.06) | 3.2 (±0.07) |
| ***Minor Alkaloids (µg/ml)*** |  |  |
| Nornicotine | 10.20 (±0.97) | 2.38 (±0.33) |
| Anatabine | 10.98 (±1.13) | 0.12 (±0.11) |
| Anabasine | 3.62 (±0.45) | 0.035 (±0.035) |
| ***Carbonyls (µg/ml)*** |  |  |
| Formaldehyde | 43.83 (±1.46) | 5.25 (±0.25) |
| Acetaldehyde | 1,691.08 (±57.02) | 3.42 (±0.29) |
| Acetone | 829.58 (±22.94) | 3.50 (±0.52) |
| Acrolein | 93.83 (±5.57.50) | ND |
| Propionaldehyde | 145.58 (±5.35) | ND |
| Crotonaldehyde | 69.25 (±3.27) | ND |
| Butanone | 268.42 (±13.19) | ND |
| Buteraldehyde | 21.25 (±1.28) | ND |
| ***Other VOCs (ng/ml)*** |  |  |
| Toluene | 489 | ND |
| Benzene | 129 | ND |
| Ethylbenzene | 87.8 | ND |
| ***ß-Carbolines (µg/ml)*** |  |  |
| Norharmane | 20.57 (±0.81) | ND |
| Harmane | 7.16 (±0.23) | ND |
| ***Solvents (mg/ml)*** |  |  |
| Propylene glycol | 1.22 (±0.15) | 12.64 (±1.6) |
| Ethylene glycol | ND | ND |

LOQ: Limit of quantitation.  ND: Not detected.

EA aerosol extracts were prepared in a similar manner. EA aerosol was generated using a Vuse Solo device containing Vuse Original (tobacco-flavored) EC liquid (4.8% nicotine concentration) in cartridges (device and cartridges purchased from vusevapor.com). A total of 800 puffs (100 puffs per filter pad) were puffed (100-ml puff volume, 4-s puff duration and 40-s interval between puffs) and final EA extracts were prepared as described above. These puffing parameters are in the range of those observed in experienced EA users (Kosmider et al.,2018). The nicotine concentration was determined for CS and EA extract as described below. Extracts were then diluted to the nicotine concentrations required for the current studies. Thus, the concentration of all constituents in extract decreased proportionally with the nicotine concentration.

Because ETOH was used as a solvent for extract preparation, both CS and EA extracts contained ≈ 7.5% ETOH after dilution depending on nicotine unit dose. To control for potential interactions between nicotine and ETOH, the same respective ETOH concentrations present in the extract doses were added to the nicotine alone doses. The 7.5% concentration of ETOH added to the highest nicotine alone dose was also added to the saline control solution. The pH of nicotine alone and extract solutions was adjusted to 7.4 with dilute NaOH, and heparin (30 units/ml) was added to help maintain catheter patency in the self-administration study. All solutions were filter-sterilized after preparation. Nicotine doses are expressed as the base. Brevital sodium (methohexital, JHP Pharmaceuticals, Rochester, MI) used for confirming catheter patency was dissolved in sterile saline.

### 2.4. Chemical analysis of extracts and nicotine solutions

#### 2.4.1. Nicotine and minor alkaloids

Nicotine and minor alkaloids samples were analyzed as described previously (Harris et al., 2022). Briefly, samples were prepared by serial dilution of the CS and EA extract with 10 mM ammonium acetate, and addition of [D3]nicotine, [D4]nornicotine, [D4]anabasineand [D4]anatabine as internal standards. Samples were then analyzed by liquid chromatography-tandem mass spectrometry (LC-MS/MS) on a Hypercarb column (Thermo Scientific), using 10 mM ammonium acetate (with 0.01% formic acid) and methanol as mobile phase.

#### 2.4.2. ß-Carbolines

Analysis of harmane and norharmane was conducted as described previously (Harris et al., 2022). CS and EA extracts (50 μl) were mixed with ^13^C2^15^N-harman and D7-norharman internal standards and diluted to 5 ml with water. The mixture was subjected to purification by solid supported liquid-liquid extraction using ChemElut cartridges (Agilent, Santa Clara,CA) and extracted using 8 ml methylene chloride twice. The samples were dried in SpeedVac for 1.5 h and reconstituted in1 ml of deionized water. Samples were then analyzed by LC-MS/MS on a Zorbax SB C18 (Agilent, Santa Clara, CA) column, using water (with 0.1% trifluoroacetic acid) and acetonitrile (with 0.1% trifluoroacetic acid) as mobile phase. The mass spectrometer with electrospray ionization source was set in the positive ion mode with selective reaction monitoring (SRM) monitoring m/z 169→115 and m/z 183→115 for norharmane and harmane, respectively.

#### 2.4.3. Carbonyl VOCs

Cigarette smoke and EA extracts were analyzed for eight carbonyl compounds as described previously (Harris et al., 2022). Briefly, CS and EA extracts (100 μl) were derivatized with 2,4-dinitrophenylhydrazine (DNPH) for 20 min. The derivatization reaction was stopped with 1% Trizma base and samples were analyzed by high-performance liquid chromatography (HPLC-UV) on a Phenomenex C18 (2) 250 mm × 4.6 mm column, using 30% acetonitrile/10%tetrahydrofuran/1% isopropanol/59% water as mobile phase A and 65% acetonitrile/1% tetrahydrofuran/1% isopropanol/33%water as mobile phase B, with the UV detector set at 365 nm.

#### 2.4.4. Other VOCs

Levels of the VOCs benzene, ethylbenzene, and toluene were measured in CS extract by the Tobacco and Volatiles Branch of the Centers for Disease Control and Prevention (Atlanta, GA) by gas chromatography-mass spectrometry (GC-MS) using standard methods described previously (Vu et al., 2021). A 0.8 ml sample of raw extract prepared as above was diluted with methanol to a final volume of 5 ml that was divided into 250 µl aliquots. Each aliquot was placed into glass vial inserts inside of an autosampler vial to minimize headspace and loss on evaporation. Vials were then stored at -80 ° C until the date of analysis. From each autosampler vial, 1-uL was injected into the GC-MS for analysis. Samples were analyzed in duplicate at one and two months after extraction.

#### 2.4.5. Glycols

Glycol concentrations were measured by gas chromatography (GC) with flame ionization detection (FID) on an Agilent 6890 GC with ChemStation, V10.1 as we have described previously (Harris et al. 2022). 1,3-propanediol (trimethylene glycol) was utilized as an internal standard. Levels greater than the limit of linearity (100 mg/dl) are diluted.

### 2.5 Procedure

#### 2.5.1. Surgical Procedures

##### Catheter implantation

Each rat was implanted with a chronic indwelling catheter into the right jugular vein under isoflurane anesthesia, described in detail elsewhere.(LeSage et al., 2003, 2002). The catheter was externalized between the scapulae and attached to the vascular-access harness (see above) that allowed connection to a fluid swivel via a tether for nicotine administration. Animals were allowed to recover for at least three days after surgery, during which time they received daily i.v. infusions of heparinized saline, ceftriaxone antibiotic (5.25 mg), and s.c. injections of buprenorphine (0.05 mg/kg; first two days only) for analgesia.

Infusions of methohexital (0.1 ml, 10 mg/ml, i.v.) were administered to check patency post-session on Fridays. If a rat failed to exhibit anesthesia within 3-5 sec during these patency checks, that animal’s data from that week were excluded from analysis. Another catheter was then implanted into a femoral vein and the rat was returned to the protocol. If that catheter failed, a third catheter was implanted into the contralateral femoral vein. If that catheter lost patency, the animal was removed from the protocol.

##### Microdialysis probe implantation

Rats used for microdialysis underwent stereotaxic surgery to implant a cranial guide cannula (Bioanalytical Systems, Inc., West Lafayette, IN) as described in detail elsewhere (Carreño et al., 2024; Villégier et al., 2007b, 2007a). Briefly, the guide cannula was implanted 2.0 mm above the target area, fixed to the skull with acrylic dental cement, and sealed with a dummy cannula. Anatomical coordinates for the NAc shell were determined from a stereotaxic atlas (Paxinos and Watson, 1989) with histological confirmation. The following guide cannula coordinates were measured from the dura; AP, + 2.0 mm; ML, ± 1.2 mm; DV, 5.8 mm.

#### 2.5.2. Drug self-administration

Four groups of rats (N=22-25 per group, 11-14 males, 11-12 females per group) were allowed to self-administer nicotine alone, CS extract, EA extract, or saline using our standard procedures (LeSage et al., 2016; LeSage et al., 2016). A nicotine unit dose of 30 µg/kg/infusion was used in the nicotine alone and extract groups. Sessions began with onset of a stimulus light above the active lever, and each press on that lever produced offset of the stimulus light and an infusion delivered in a volume of 0.1 ml/kg in approximately 1 sec. Each infusion was immediately followed by a 15-sec timeout during which the stimulus light remained off and lever presses were recorded but had no programmed consequence. Following the timeout, the stimulus light was illuminated to indicate drug availability. Presses on the inactive lever were recorded but had no programmed consequence. The active lever was baited with food powder on the first session only to ensure contact with the reinforcement contingency. Data from this session were not analyzed. Sessions were 2 hr in duration and typically ran five days per week, depending on national holidays. Infusions were initially available under a fixed-ratio (FR) 1 schedule for at least 10 sessions and until at least five infusions were earned per session and no significant trend in the number of infusions was apparent across five sessions. The FR was increased to FR 2 for at least 5 sessions and until responding was stable (no trend in infusions for five consecutive sessions), and then to FR 3 for at least 10 sessions until responding was stable. Rats were considered to have acquired self-administration at FR 3 if they earned at least five infusions per session, showed a mean active:inactive response ratio of at least 2:1, and no significant trend in responding across three consecutive sessions.

Rats in the nicotine alone and extract groups that acquired self-administration at FR 3 (14-16 total, 9 males, 5-7 females per group) were used to assess behavioral economic demand for nicotine. During this phase, unit price was increased weekly (Mondays) by reducing the nicotine dose to 15, 7.5, 4, 2, 1, and 0 µg/kg/infusion (saline). This progression is similar to that used in our previous studies using a unit dose-reduction protocol to measure elasticity of demand, and the dose range is broad enough to yield a bi-tonic dose response curve (Grebenstein et al., 2013, 2015). Some rats in the saline group met acquisition criteria, suggesting that the visual stimuli and/or ETOH may have had weak reinforcing effects. Those rats were exposed to weekly reductions in ETOH concentration to assess whether ETOH was reinforcing. ETOH concentrations were matched to the concentrations in the nicotine alone and extract groups during nicotine dose reduction (3.8, 1.9, 0.95, 0.48, 0.24, and 0%).

Following behavioral economic demand assessment, nicotine and extract rats were allowed to reacquire stable SA and then placed on extinction by substituting saline for the training formulation for at least 10 sessions and until extinction criteria were met (greater than 50% reduction and no significant trend in infusions earned per session over three consecutive sessions). Stress-induced reinstatement was then measured on Tuesday and Friday sessions provided extinction criteria were met during intervening sessions. During reinstatement tests, rats were injected i.p. with the alpha-2 adrenergic antagonist yohimbine (0, 1.25, or 2.5 mg/kg) 30 min prior to the session. At these doses, yohimbine acts as a pharmacological stressor and can increase plasma stress hormone (corticosterone) levels and elicit reinstatement of extinguished nicotine/extract-seeking behavior in rats (Costello et al., 2014; Gellner et al., 2023).

#### 2.5.3. In Vivo microdialysis

Procedures were similar to our routine approach (Carreño et al., 2024; Villégier et al., 2007b, 2007a). Rats were implanted with an i.v. catheter and microdialysis guide cannula aimed at the shell of the NAc as described above. Animals were given 2 days to recover with daily handling after surgeries. Catheters were flushed daily, and animals were habituated to the micro-dialysis chambers for 5 min per day. Intravenous catheter patency was tested by 0.2 ml propofol 1 day before and immediately after the experiment. On the experimental day, the dummy cannula was replaced with a 2-mm microdialysis probe and microdialysis was carried out under free-moving conditions using our standard methods. After a 4-h equilibration period, rats (N=4-7/group) received two i.v. infusions of saline, CS extract, EA extract, or nicotine alone via the catheter with a 1 min interinfusion interval. The nicotine dose for each nicotine formulation was 30 μg/kg/inf. Samples were then collected via the microdialysis probe every 20 min during 1 h. DA levels in samples were measured using high performance liquid chromatography with electrochemical detection using our routine procedures. The placement of microdialysis probes in the NAc was verified histologically.

#### 2.5.4. In vitro binding and function at nAChRs

Binding assays of nAChRs to [^3^H]-epibatidine were conducted with nicotine alone and extracts using human embryonic kidney (HEK) 293 cell lines stably expressing human α4ß2 and α3ß4 nAChR subtypes, or the rat α7 nAChR subtype. Assays were also conducted using α4ß2 or α7 nAChRs expressed in rat forebrain or cortex, respectively. Binding assays were conducted in three (α4β2, α7) or four (α3β4) independent experiments, with primary screening assays performed in quadruplicate and secondary competition curves generated from 10 concentrations in singlet. Agonist and antagonist function of these nAChR subtypes were measured using an aequorin bioluminescence assay (Karadsheh et al., 2004). Functional aequorin assays were conducted in 7-9 (α4β2) or 4-5 (α3β4) independent experiments per formulation. Cell lines transfected to express the bioluminescent aequorin protein were incubated for 3 hours with coelenterazine, which complexes with aequorin in the cell. When the complex binds calcium, it emits light that is directly proportional to the calcium concentration inside the cell that results from the opening of the receptor ion channel. Agonist activity of a compound is measured by quantifying the light it induces on a luminescence plate reader, while antagonist activity is measured by blockade of light induced by 10 μM nicotine.

#### 2.5.5. In Vitro binding and function at non-nAChRs

Binding and functional activity of a single sample of nicotine alone and extracts were measured at a wide range of CNS and enzyme-linked receptor targets by Eurofins Discovery (San Diego, CA) using their DiscoverX SAFETYscan E/IC50 ELECT commercial service. Assays utilized PathHunter enzyme fragment complementation (EFC) technology, FLIPR®-based cellular screening assays, KINOMEscan kinase binding assays, calcium fluorescence assays, cAMP second messenger assays, and a variety of standard enzymatic assays.

Supplementary Table 1 lists all of the targets and associated control compounds. Details of the analyses are available from the Eurofins Discovery website (https://www.eurofinsdiscovery.com/4567d9bd-97f8-4ef2-a669-5b6660f43eb8). Binding analysis was initially done for each formulation at a high nicotine concentration (0.03 mg/ml, 184.9 µM) to exploit the potential for binding by non-nicotine constituents in extracts, which are present at much lower concentrations than nicotine, and determine whether functional analysis was warranted. Functional analyses of nicotine alone and CS extract were done using a broad range of nicotine concentrations that included those estimated in brains of smokers (0.36 -1.4 µM) (Rose et al., 2010a) and in rats self-administering nicotine at the training dose used in the study (30 µg/kg/inf **≈** 1.2 µM, (Donny et al., 2000). Because binding data showed no significant differences between nicotine alone and EA extract at any receptor (see below), EA extract was not examined for functional activity.

### 2.6. Data analysis

#### 2.6.1. Self-administration

##### Dependent measures

Mean lever presses on the active and inactive lever, number of infusions, and nicotine intake across the last three sessions at each nicotine unit dose were primary dependent measures. Dose-response data were also expressed as % baseline (30 µg/kg) to control for the baseline difference between nicotine alone and CS extract (Fig 1A and 3A) and directly measure the magnitude of compensation in each rat. The reinforcement threshold for each formulation was defined as the lowest unit dose that differed significantly from 0 µg/kg. In addition, elasticity of demand served as the primary measure of relative reinforcing efficacy, which was determined by exponential demand curve analysis using the following equation:^26^

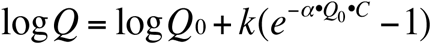

**Figure 1.**
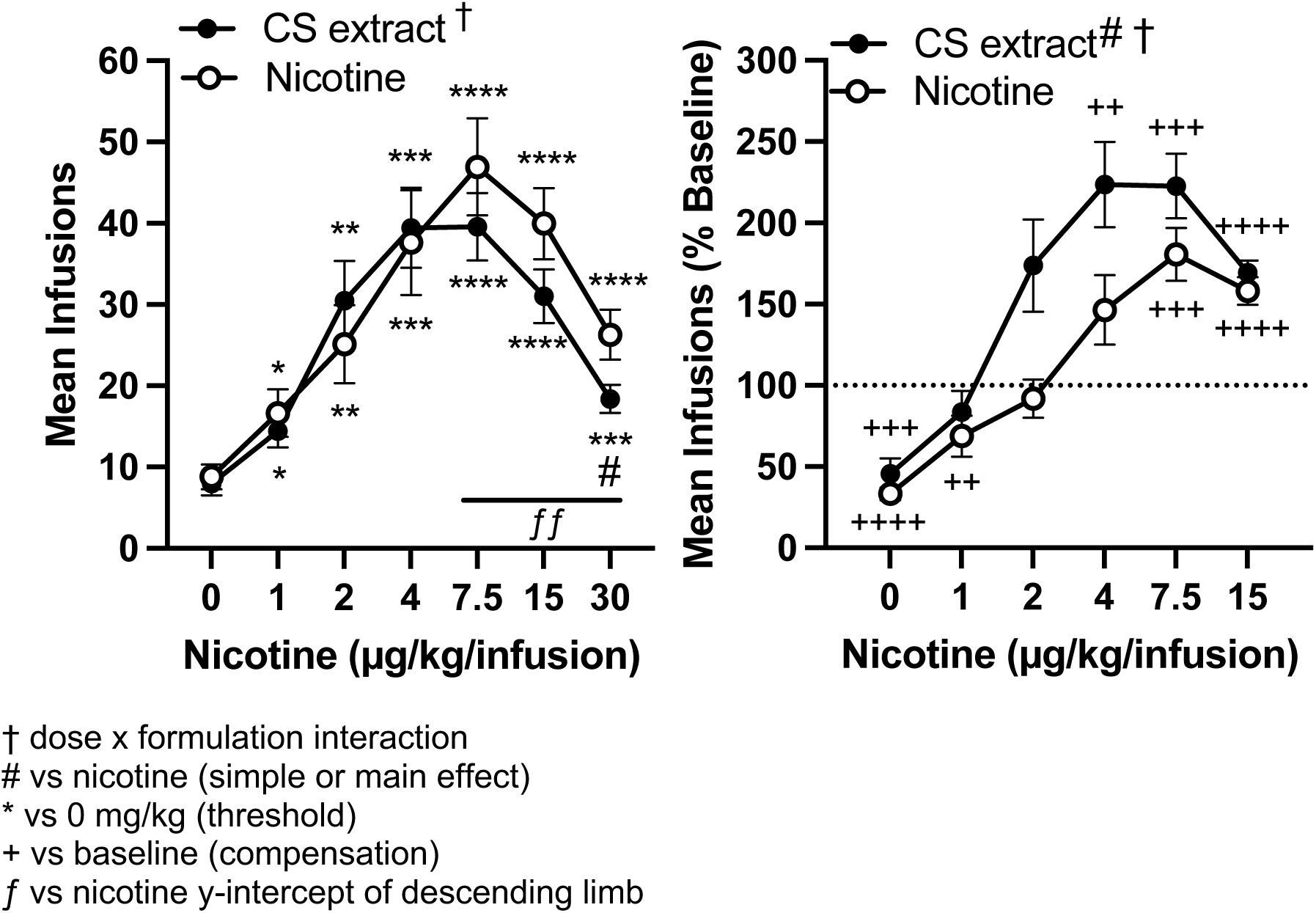
Mean (±SEM) number of infusions during the course of unit dose reduction expressed as absolute values (left panel) or a percentage of baseline (30 µg/kg, right panel) in rats responding for nicotine alone or CS extract at equivalent nicotine doses. Left panel: Significant differences from saline, *p < 0.05, **p < 0.01, ***p < 0.001, ****p < 0.0001. Significant unit dose x formulation interaction, † p < 0.05. Significant difference from CS extract # p < 0.05. Significant difference in y-intercept across 7.5 to 30 µg/kg, ƒƒ p < 0.01. Right panel: Significant difference from baseline, ++ p < 0.01, +++ p < 0.001, ++++ p < 0.0001. Main effect of formulation, # p < 0.05. Significant unit dose x formulation interaction, † p < 0.05.

The dependent variable, *Q*, is the quantity consumed. The independent variable, *C*, is the cost of nicotine based on the unit price (FR/unit dose). The free parameters, *Q*_0_ and *α* are estimated from the best-fit function and refer to the maximum level of consumption at zero price (i.e., level or “intensity” of demand) and the rate of change in consumption with increases in unit price, respectively. The range of the exponential function, *k*, is a constant specifying the range of consumption in log units. The *k* value is held constant across all data sets being compared (set to 2.401 in the present study), because changes in *k* impact the value of *α*. The *α* parameter is considered a measure of reinforcing efficacy, such that drugs that produce rapidly declining (elastic) demand curves have higher *α* values and lower reinforcing efficacy than drugs that produce slower declining (inelastic) demand curves. Therefore, *α* served as the index of elasticity of demand for, or reinforcing efficacy of, nicotine and extracts. Other demand measures of interest included: *Q*_0_, the level or intensity of demand as described above; Omax, the maximal level of active lever responses; and Pmax, the unit price (responses per unit dose) at which Omax occurs and consumption becomes relatively elastic. Omax and Pmax were observed values, not estimates. Demand functions were generated using a template for GraphPad Prism software (GraphPad Software, Inc; La Jolla, CA) provided by the Institutes for Behavior Resources, Inc. (Baltimore, MD) on their website (http://www.ibrinc.org/index.php?id=181). The mean of parameter values fit to individual rat data served as the primary measures of demand. Finally, a reinstatement score (mean active lever - inactive lever presses during reinstatement tests) was used as the measure of yohimbine-induced reinstatement to control for non-specific motor effects of yohimbine observed in some rats.

##### Statistical analyses

Stable baseline intake (mean of the last three sessions of acquisition at the 30 µg/kg dose) was compared between each formulation and nicotine alone using two-way ANOVA (formulation x sex). Dose-response data during dose reduction was compared between each extract and nicotine alone using a three-factor (sex x dose x formulation) mixed effects model with the maximum likelihood method and Geisser-Greenhouse correction, followed by Dunnett or Bonferroni tests for within-or between-group multiple comparisons, respectively. Separate two-factor (dose x formulation) mixed-effects models were similarly conducted to examine effects within each sex and sex effects within each formulation (dose x sex). In light of an apparent shift in the descending portion of the dose-response curve (7.5 - 30 µg/kg) for CS and EA extract versus nicotine alone, where non-nicotine constitutents are less diluted, the slopes and y-intercepts of absolute infusion rates over this dose range were compared via linear regression. A reduction in the y-intercept of the descending limb of the dose-response curve with no change in slope (lines are parallel) also indicates a reduction in the x-intercept, reflecting a leftward shift of the curve. Mean demand parameters were compared between each extract and nicotine alone using independent-samples *t*-tests with Welch’s correction. Pmax values were not normally distributed and no type of transformation fixed this, so these values were compared via nonparametric Mann-Whitney U tests.

Reinstatement scores were compared between each extract and nicotine alone using a three-factor (sex x dose x formulation) mixed effects model with the maximum likelihood method and Geisser-Greenhouse correction, followed by Dunnett or Bonferroni tests for within- or between-group multiple comparisons, respectively. Residuals were not normally distributed for the analysis on absolute scores, so it was done using log transformed data.

#### 2.6.2. Microdialysis

Data were analyzed using two-way analysis of variance, with formulation as a between-subjects factor and time as a within-subjects factor, followed by Bonferroni- or Dunnett’s-post-hoc comparisons. Data were analyzed with JMP statistical analysis software (SAS Institute, Cary, NC).

#### 2.6.3. Receptor binding and function

Data for binding affinity and functional activity at nAChRs were analyzed by nonlinear least-squares regression to obtain Ki or EC50 values, respectively (see http://pdsp.med.unc.edu for further details). For functional activity at non-nAChRs. data were analyzed by non-linear least-squares regression with the Levenberg-Marquardt algorithm. This procedure only attempts to fit a curve and calculate EC50 values if the maximum response at the highest concentration is above 50%. Statistically significant differences in Ki or EC50 between nicotine alone and each extract were determined via unpaired Welch’s-corrected t-tests.

## 3. Results

### 3.1. Self-administration dose-response curves for CS extract versus nicotine alone

#### 3.1.1. Absolute infusion rates (Figure 1 left panel)

Infusion rates during acquisition <u>in</u> <u>all rats</u> were higher for nicotine alone and CS extract compared to saline (Supplementary Fig 1). There was a tendency for infusion rates to be lower for CS extract than nicotine alone, but this was not significant (Supplementary Fig 1). In addition, CS extract did not differ from nicotine alone in terms of the number of sessions to or proportion of rats to meet criteria (not shown). However, baseline rate at the end of acquisition (30 µg/kg) <u>in rats that acquired</u> self-administration was significantly lower for CS extract compared to nicotine alone (F = 5.63, p<0.05). During dose reduction, there was no main effect of formulation, but there was a significant main effect of dose (F=45.77, p<0.0001) and sex (F=8.54, p<0.01), and dose x sex (F=2.87, p<0.05) and dose x formulation (F=2.31, p<0.05) interactions. The dose response curve was an inverted U-shape for both formulations, with initial compensatory increases in infusion rate as the unit dose was decreased to a peak rate at 7.5 µg/kg, followed by decreases in infusion rate at lower doses. The slope of the descending limb of the dose-response curve (7.5 - 30 µg/kg) did not differ between CS extract and nicotine alone, but the y-intercept of the curve was significantly lower for CS extract versus nicotine alone (45.9 ± 3.9 vs 53.8 ± 5.5, respectively, F = 5.9, p < 0.05), indicating a downward/leftward shift) in this portion of the curve for CS extract relative to nicotine alone. There were no differences in mean infusions across the ascending limb of the dose-response curve, with a reinforcement threshold of 1.0 µg/kg for both CS extract (t=3.25, p<0.05) and nicotine alone (t=3.34, p<0.05). Thus, potency did not differ between formulations in the ascending portion of the curves. The ratio of active:inactive responding was greater than 2:1 at all unit doses (Supplementary Figure 2).

##### Infusion rates as a percentage of baseline (Figure 1, right panel)

There was a main effect of formulation (F=5.57, p<0.05), dose (F=42.83, p<0.0001), and dose x formulation interaction (F=3.46, p<0.01) on infusion rate as a percentage of baseline, but no effect of or interactions with sex. Although there were no significant differences between formulations at each nicotine dose, the dose-response curve was shifted upward/leftward for CS extract compared to nicotine alone, indicating a greater overall compensatory increase in infusion rate for CS extract. Moreover, significant compensatory increases in infusion rate during dose reduction occurred over a wider range of unit doses for CS extract (4 - 15 µg/kg) compared to nicotine alone (7.5 - 15 µg/kg). Infusion rates were significantly below baseline at the 1 µg/kg unit dose in the nicotine alone group (q=4.01, p<0.01), but not the CS extract group, indicating greater persistence of responding during dose-reduction in the CS extract group.

#### 3.1.2. Sex differences (Figure 2)

Follow-up two-factor analyses indicated there was a main effect of sex (F=8.98, p<0.01) and formulation (F=5.63, p<0.05) but no sex x formulation interaction on absolute baseline infusion rate at the end of acquisition (30 µg/kg), with higher infusion rates in females than males and lower infusion rates for CS extract than nicotine alone. Multiple comparisons showed infusion rates were higher in females than males (t=3.01, p<0.01) for nicotine alone, but not for CS extract. Infusion rates for CS were lower than nicotine alone in females (t=2.19, p<0.05), but not males. During dose reduction, overall absolute infusion rates averaged across formulations were higher in females compared to males (F=9.79, p<0.01). Within each formulation, absolute infusion rates for nicotine alone, but not CS extract, were significantly higher in females compared to males (F=5.43, p<0.05). Among males, there was a significant effect of dose (F=25.97, p<0.0001) on absolute infusion rates, but no effect of formulation or dose x formulation interaction. However, the dose-response analysis within each formulation for males indicated that the reinforcement threshold was lower for CS extract (2.0 µg/kg, q=5.26, p<0.01) than nicotine alone (4.0 µg/kg, q=5.29, p<0.01). Among females, there was a significant effect of dose (F=19.68, p<0.0001) on absolute infusion rates, but no effect of formulation or dose x formulation interaction. However, dose-response analysis within each formulation indicated that the reinforcement threshold was lower for CS extract (4.0 µg/kg, q=4.77, p<0.05) versus nicotine alone (7.5 µg/kg, q=6.39, p<0.01). The slope and y-intercept of the descending limb of the dose-response curve (7.5 - 30 µg/kg) did not differ between CS extract and nicotine alone for either sex, but the lower y-intercept for CS extract versus nicotine alone approached significance in females (p = 0.06). As noted above, there were no sex differences in infusion rates as a percentage of baseline.

**Figure 2.**
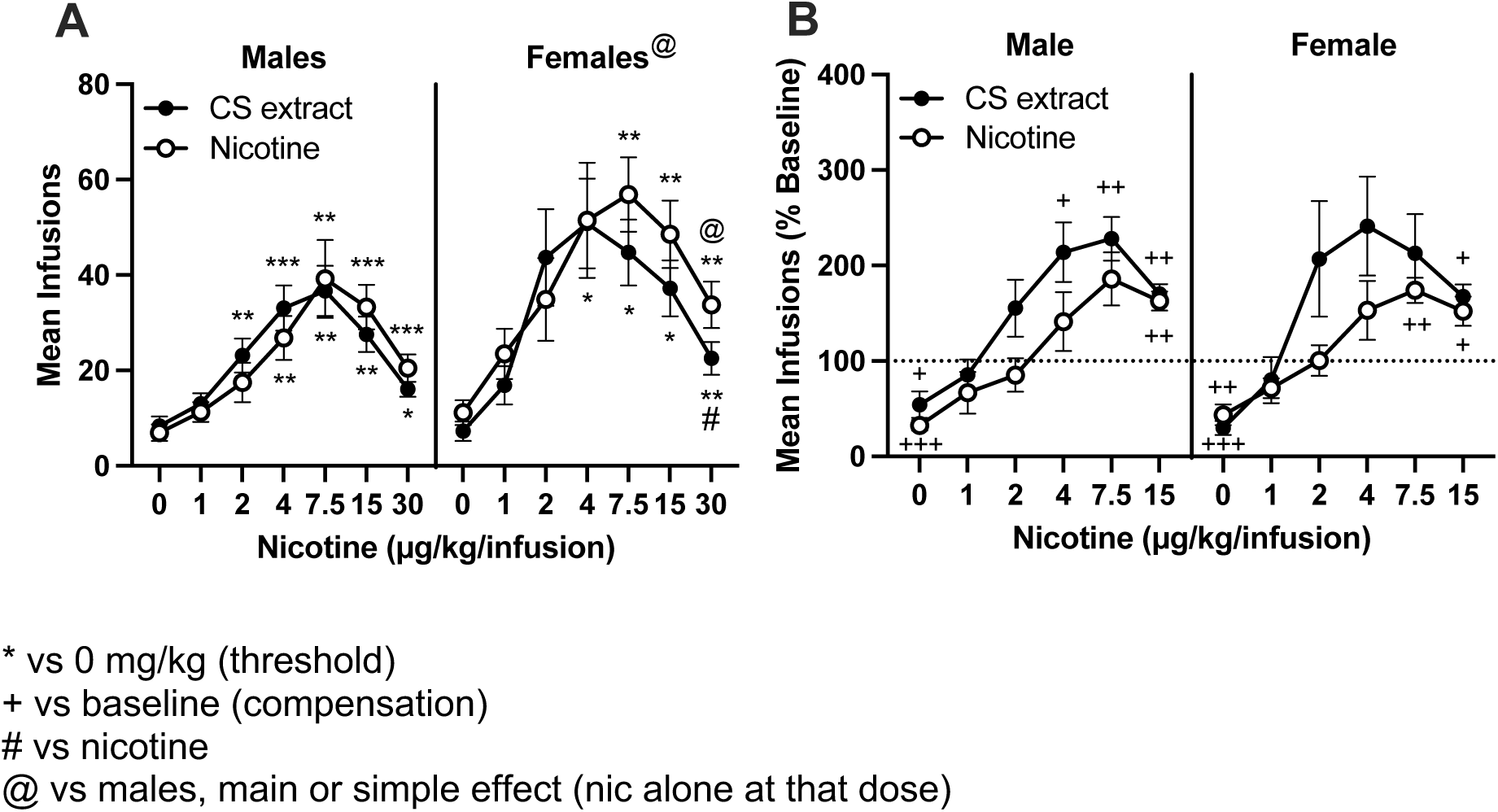
Mean (±SEM) number of infusions during the course of unit dose reduction expressed as absolute values (**A**) or a percentage of baseline (30 µg/kg, **B**) in male (left panels) and female (right panels) rats responding for nicotine alone or CS extract at equivalent nicotine doses. **A:** Significant differences from saline, *p < 0.05, **p < 0.01, ***p < 0.001. Significant difference from CS extract # p < 0.05. Significant main effect of sex or different from males at that nicotine alone unit dose, @ p < 0.05. **B:** Significant difference from baseline, ++ p < 0.01, +++ p < 0.001, ++++ p < 0.0001. Main effect of formulation, # p < 0.05. Significant unit dose x formulation interaction, † p < 0.05.

### 3.2. Self-administration dose-response curves for EA extract versus nicotine alone

#### 3.2.1. Overall absolute infusion rates (Figure 3, left panel)

Infusion rates during acquisition were higher for nicotine alone and EA extract compared to saline (Supplementary Fig 1). Infusion rates in all rats during acquisition for EA extract appeared lowed compared to nicotine alone, but this apparent difference was not significant (Supplementary Fig 1). In addition, EA extract did not differ from nicotine alone in terms of sessions to meet acquisition criteria or the proportion of rats meeting those criteria (not shown). Absolute baseline infusion rate at the end of acquisition (30 µg/kg) in rats that acquired self-administration was lower for EA extract versus nicotine alone (F=4.68, p<0.05). During dose reduction, there was a significant main effect of dose (F=37.5, p<0.0001) and sex (F=5.5, p<0.05), but the effect of formulation only approached significance (p<0.1) and no interactions were significant. The shape and position of the dose-response curve were similar between formulations, with infusion rates increasing to a peak at 7.5 µg/kg followed by decreases in infusion rate at lower doses. Although the curve for EA extract appeared lower compared to nicotine alone, none of the differences at each dose were significant. The slope of the descending limb of the dose-response curve (7.5 -30 µg/kg) did not differ between EA extract and nicotine alone, but the y-intercept of the curve was significantly lower for EA extract versus nicotine alone (39.01 ± 4.1 vs 53.79 ± 5.6, respectively, F = 9.8, p < 0.01), indicating a downward/leftward shift in this portion of the curve for EA extract relative to nicotine alone. The ascending limb was similar and the reinforcement threshold was 1 µg/kg for both formulations. The ratio of active:inactive responding was greater than 2:1 at all unit doses (Supplementary Figure 2).

**Figure 3.**
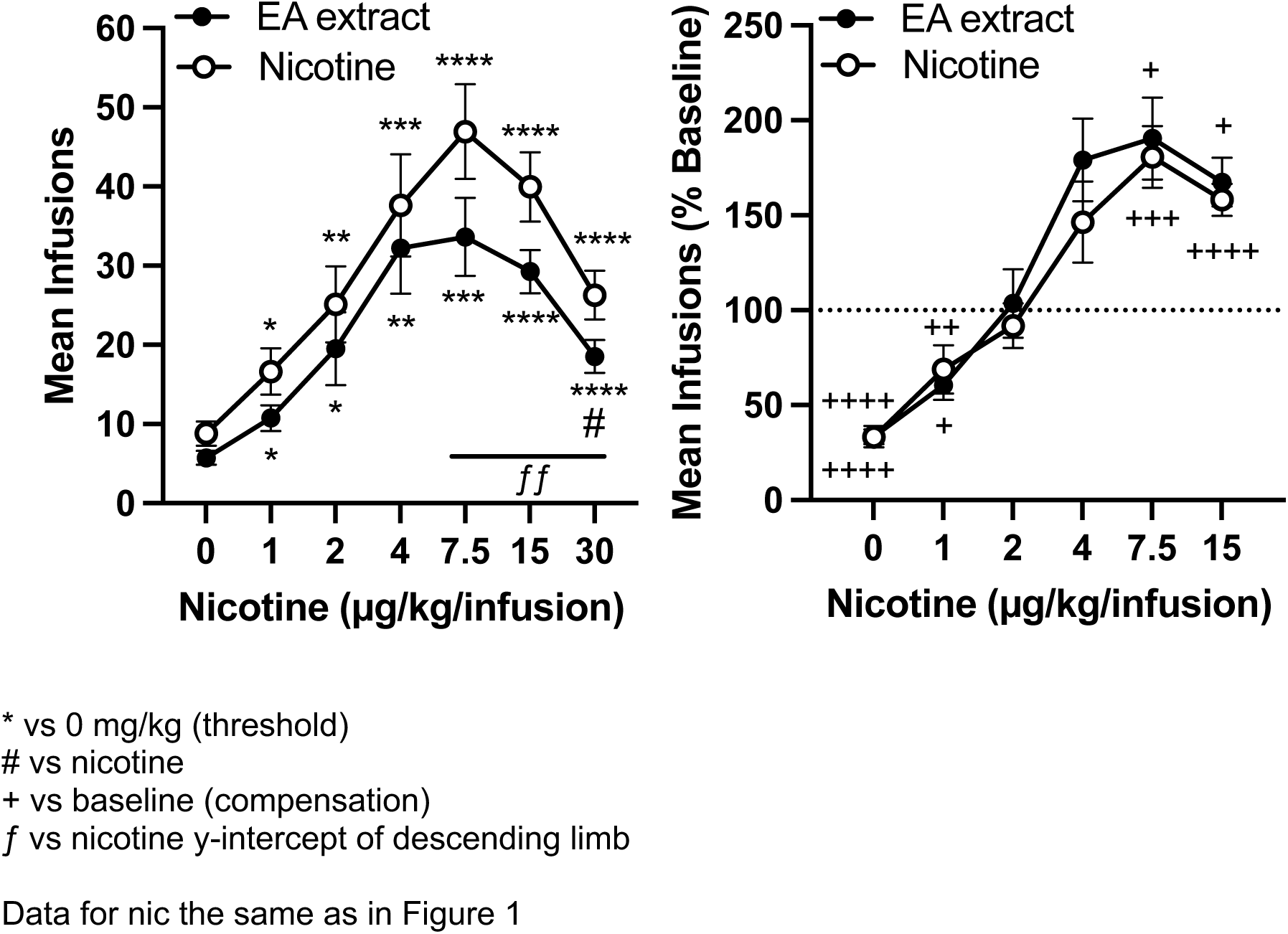
Mean (±SEM) number of infusions during the course of unit dose reduction expressed as absolute values (left panel) or a percentage of baseline (30 µg/kg, right panel) in rats responding for nicotine alone or EA extract at equivalent nicotine doses. Data for nicotine alone are the same as in Figure 1. Left panel: Significant differences from saline, *p < 0.05, **p < 0.01, ***p < 0.001, ****p < 0.0001. Significant difference from EA extract # p < 0.05. Significant difference in y-intercept across 7.5 to 30 µg/kg, ƒƒ p < 0.01. Right panel: Significant difference from baseline, ++ p < 0.01, +++ p < 0.001, ++++ p < 0.0001.

##### Infusion rates as a percentage of baseline (Figure 3, right panel)

There was a main effect of dose (F=45.12, p<0.0001), but no other effects or interactions on infusion rates as a percentage of baseline. The dose-response curves were similar in shape and relative position between formulations, and compensatory increases in infusion rate during dose reduction occurred over the same unit dose range for EA extract and nicotine alone (7.5 - 15 µg/kg).

#### 3.2.2. Sex differences, absolute infusion rates (Figure 4A)

Follow-up two-factor analysis indicated there was a main effect of sex (F=10.49, p<0.01) and formulation (F=4.68, p<0.05) but no sex x formulation interaction on absolute baseline infusion rate at the end of acquisition (30 µg/kg). Multiple comparisons showed infusion rates were higher in females than males (F=2.92, p<0.01) for nicotine alone, but not for EA extract. During dose reduction, overall absolute infusion rates averaged across formulations were higher in females compared to males (F=4.34, p<0.05). Within each formulation, absolute infusion rates for nicotine alone were higher in females (F=5.43, p<0.05), but there was no sex difference for EA extract. Among males, there was a significant effect of dose (F=29.18, p<0.0001) on absolute infusion rates, but no effect of formulation or dose x formulation interaction. Dose-response analyses within each formulation in males indicated that the reinforcement threshold was lower for EA extract (1.0 µg/kg, q=4.34, p<0.05) than nicotine alone (4.0 µg/kg, q=5.29, p<0.05). Among females, there was a significant effect of dose (F=12.73, p<0.001) on absolute infusion rates, but no effect of formulation or dose x formulation interaction. Dose-response analyses within each formulation indicated that the reinforcement threshold was 30 µg/kg for EA extract (q=4.47, p<0.05) and 7.5 µg/kg for nicotine alone (q=6.39, p<0.01), with relatively high variability in EA extract SA on the ascending limb of the dose-response curve compared to nicotine alone. The slope of the descending limb of the dose-response curve did not differ between EA extract and nicotine alone for either sex. However, the y-intercept for EA extract was lower versus nicotine alone in both males (37.6 ± 3.1 vs 49.8 ± 6.5, respectively, F = 4.9, p < 0.05) and females (48.0 ± 11.6 vs 64.3 ± 8.0, respectively, F = 5.3, p < 0.05), reflecting a downward/leftward shift similar to Figure 3 (left).

**Figure 4.**
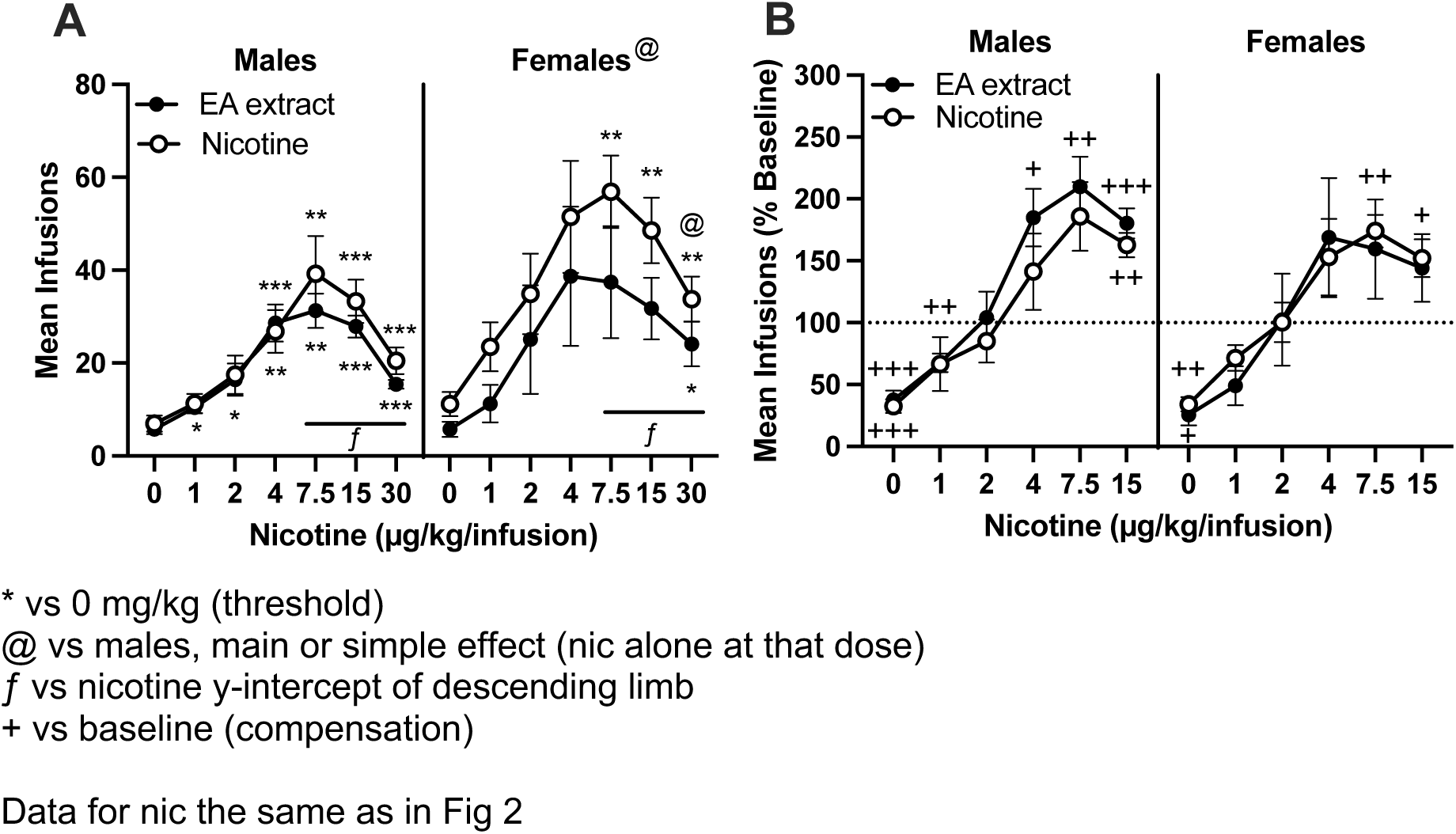
Mean (±SEM) number of infusions during the course of unit dose reduction expressed as absolute values (**A**) or a percentage of baseline (30 µg/kg, **B**) in male (left panels) and female (right panels) rats responding for nicotine alone or EA extract at equivalent nicotine doses. Data for nicotine alone are the same as in Figure 2. **A:** Significant differences from saline, *p < 0.05, **p < 0.01, ***p < 0.001. Significant main effect of sex or different from males at that nicotine alone unit dose, @ p < 0.05. Significant difference in y-intercept across 7.5 to 30 µg/kg, ƒ p < 0.05. **B:** Significant difference from baseline, + p < 0.05, ++ p < 0.01, +++ p < 0.001.

##### Infusion rates as a percentage of baseline (Figure 4B)

There was no sex difference overall or within each formulation. The dose-response curve for EA extract as a percentage of baseline was generally similar to nicotine alone in each sex. There was a significant effect of dose for males (F=30.2, p<0.0001) and females (F=18.7, p<0.0001), but no effect of formulation or dose x formulation interaction. However, compensatory increases in infusion rate occurred over a wider unit dose range for EA extract (4 -15 µg/kg) versus nicotine alone (15 µg/kg) in males, while the opposite occurred in females (no compensation for EA extract, compensation at 7.5 - 15 µg/kg for nicotine alone).

### 3.3. Behavioral economic demand analysis

Table 2 shows parameter estimates from the exponential demand curve analysis in each sex and overall. There were no sex differences in any of the parameters. Overall, Q0 (intensity of demand) was significantly lower for CS extract compared to nicotine alone (t = 2.84, p < 0.05), while the difference in Q0 between EA extract and nicotine alone approached significance (p=0.09). The higher Pmax for CS extract compared to nicotine alone was nearly significant (U = 67.0, p = 0.05). There were no differences in any other parameters between either extract and nicotine alone.

**Table 2.** Mean (±SEM) exponential demand curve parameters for each formulation in each sex and overall (*k* = 2.401).

| **Formulation** | **α** | **Q<sub>0</sub>** | **P<sub>max</sub>** | **O<sub>max</sub>** | **r<sup>2</sup>** |
|---|---|---|---|---|---|
| **Males** |  |  |  |  |  |
| Nicotine alone | 0.000795 (0.000118) | 1.07 (0.17) | 516.7 (133.9) | 132.8 (20.3) | 0.96 (0.016) |
| CS extract | 0.000733 (0.000069) | 0.67 (0.09) | 761.1 (149.7) | 124.2 (14.4) | 0.97 (0.009) |
| EA extract | 0.000818 (0.000084) | 0.80 (0.12) | 572.2 (73.18) | 108.6 (9.3) | 0.97 (0.013) |
| **Females** |  |  |  |  |  |
| Nicotine alone | 0.000488 (0.000109) | 1.38 (0.27) | 550.0 (172.9) | 183.0 (27.1) | 0.97 (0.007) |
| CS extract | 0.000530 (0.000090) | 0.84 (0.11) | 790.0 (207.0) | 168.1 (21.8) | 0.98 (0.007) |
| EA extract | 0.000640 (0.000153) | 1.00 (0.18) | 708.3 (191.2) | 187.6 (58.9) | 0.97 (0.006) |
| **Overall** |  |  |  |  |  |
| Nicotine alone | 0.000661 (0.000089) | 1.21 (0.15) | 531.3 (103.1) | 154.8 (17.1) | 0.97 (0.009) |
| CS extract | 0.000660 (0.000059) | **0.73 (0.07)\*** | 771.4 (116.6)^ | 139.9 (13.0) | 0.97 (0.006) |
| EA extract | 0.000747 (0.000079) | 0.88 (0.10) | 626.7 (85.9) | 140.2 (25.1) | 0.97 (0.008) |

Different from nicotine alone, \*p < 0.05,  ^p = 0.05.

### 3.4. Stress-induced reinstatement

There were no differences between nicotine alone and either extract in infusion rates or number of sessions to meet criteria during extinction (Supplementary Figure 3). There were no sex differences in reinstatement scores between either extract and nicotine alone. For CS extract (Figure 5, left panel), there was a main effect of yohimbine dose (F=21.11, p<0.0001), but no effect of formulation or dose x formulation interaction. Both doses of yohimbine significantly increased responding in both groups. For EA extract, there was a main effect of yohimbine dose (F=19.05, p<0.0001) and formulation (F=5.69, p<0.05), but no significant interaction. Although both doses of yohimbine reinstated responding in EA extract rats, the magnitude of reinstatement was weaker at 1.25 mg/kg yohimbine for EA extract compared to nicotine alone rats (t=3.09, p<0.05).

**Figure 5.**
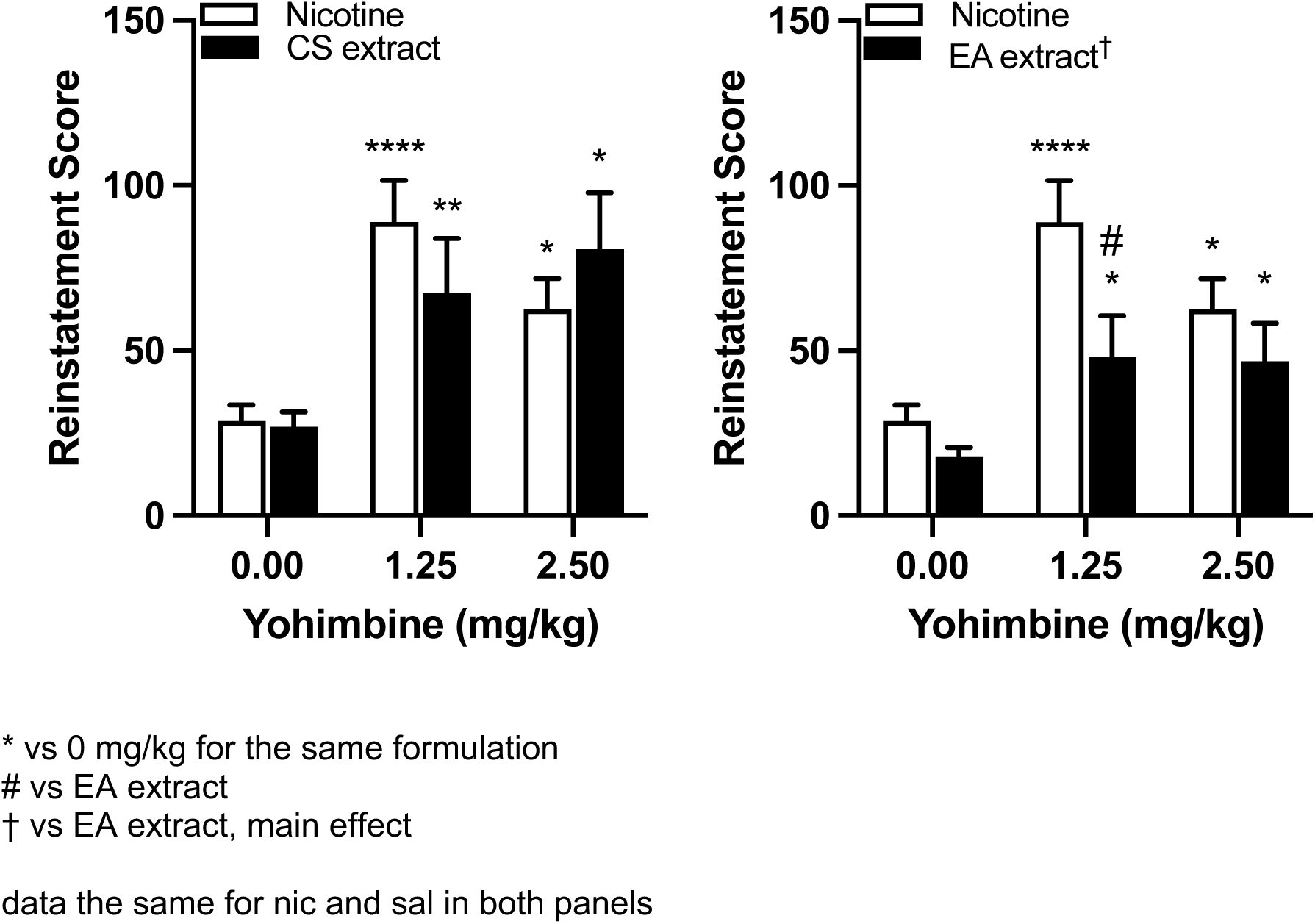
Mean (±SEM) reinstatement scores (active -inactive responses) in rats previously responding for CS extract (left panel) or EA extract (right panel) compared to nicotine alone during yohimbine primed reinstatement tests. Significant differences from saline, *p < 0.05, **p < 0.01, ****p < 0.0001. Significant difference from nicotine alone, # p < 0.05.

### 3.5. SA of saline

Seven rats (four male, three female) in the saline group met acquisition criteria at FR 3, but their responding did not vary with reductions in ETOH concentration (Supplementary Fig 4).

### 3.6. DA levels in NAc shell

Figure 6 shows extracellular levels of DA in nucleus accumbens shell as a percentage of baseline at 20-60 min after i.v. infusions of each formulation. The was significant formulation x time interaction (F=3.78, p<0.05) but no main effect of formulation or time. Nicotine alone, CS extract, and EA extract increased DA levels compared to saline at 20 min post infusion (p<0.05), but not 40 or 60 min. DA levels were generally more variable for each extract compared to nicotine alone (CS, F=4.01, p<0.01; EA, F=3.63, p<0.01).

**Figure 6.**
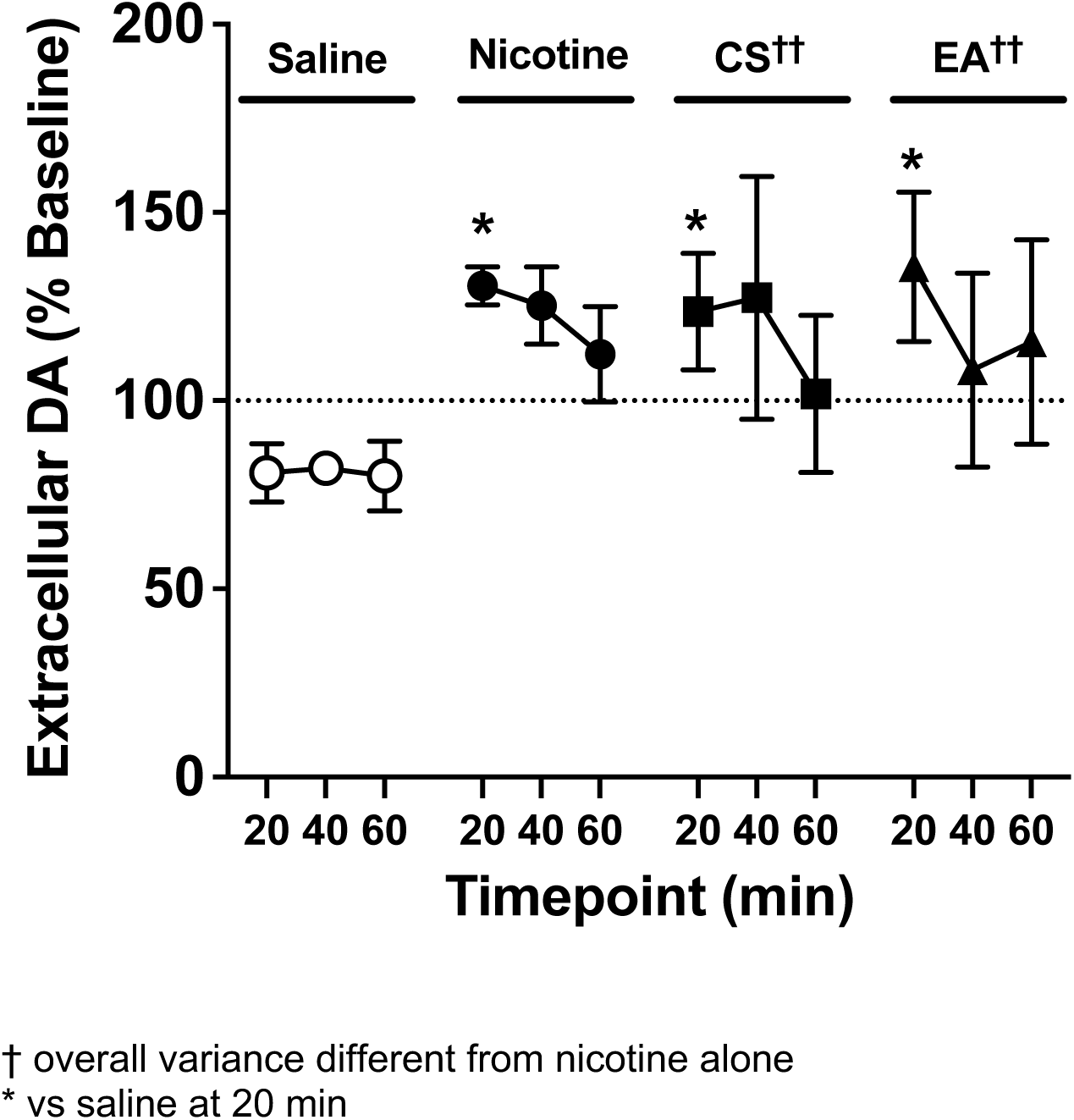
Mean (±SEM) extracellular DA levels in nucleus accumbens shell as a percentage of baseline at 20, 40, and 60 min post injection of saline, nicotine alone, CS extract, or EA extract at a nicotine dose of 0.03 mg/kg/infusion x 2, 1 min apart. Different from saline at that time point, *p < 0.05. Difference in overall variance from nicotine, †† p < 0.01.

### 3.7. Binding and function at nAChRs

Table 3 shows Ki values for competitive binding against [^3^H]-epibatidine by nicotine alone and each extract at α4ß2, α3ß4, and α7 nAChRs expressed in HEK cells or rat brain. There were no significant differences in Ki values between formulations. Also shown in Table 3 are EC50 values for luminescence intensity (% maximum relative light units (RLU)) stimulated by nicotine alone and each extract at α4ß2 or α3ß4 nAChRs expressed in HEK cells. The EC50 for CS extract was significantly lower (i.e., functional effects were significantly greater) compared to nicotine alone at α4ß2 nAChRs (t = 2.46, p < 0.05). No other differences were observed.

**Table 3.**
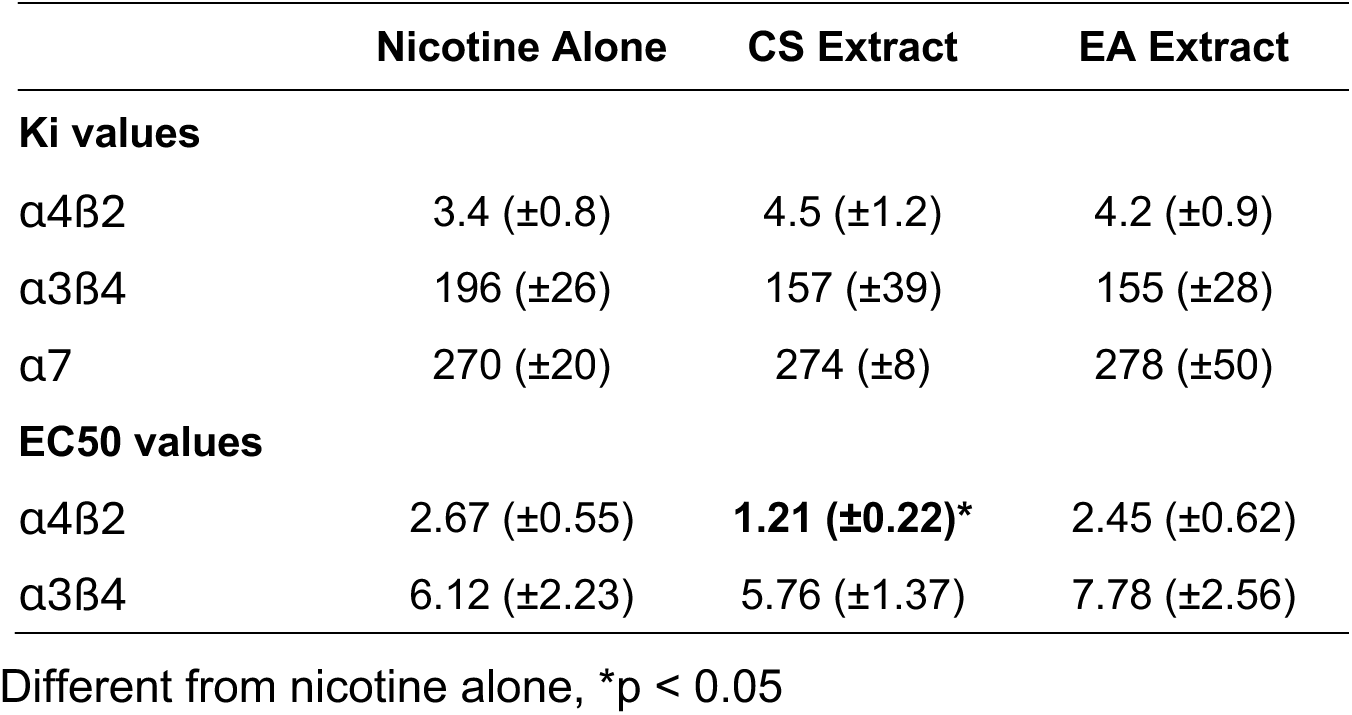
Mean (±SEM) Ki values for binding by nicotine alone and each extract at α4ß2, α3ß4, and α7 nAChRs expressed in HEK cells or rat brain; and mean (±SEM) EC50 values for agonist activity (% maximum RLU) during functional testing of formulations at α4ß2 or α3ß4 nAChRs expressed in HEK cells.

|  | **Nicotine Alone** | **CS Extract** | **EA Extract** |
|---|---|---|---|
| **Ki values** |  |  |  |
| α4ß2 | 3.4 (±0.8) | 4.5 (±1.2) | 4.2 (±0.9) |
| α3ß4 | 196 (±26) | 157 (±39) | 155 (±28) |
| α7 | 270 (±20) | 274 (±8) | 278 (±50) |
| **EC50 values** |  |  |  |
| α4ß2 | 2.67 (±0.55) | **1.21 (±0.22)\*** | 2.45 (±0.62) |
| α3ß4 | 6.12 (±2.23) | 5.76 (±1.37) | 7.78 (±2.56) |

Different from nicotine alone, \*p < 0.05

### 3.8. Binding and function at non-nAChRs

Table 4 shows % inhibition by nicotine alone and each extract (0.03 mg/ml nicotine concentration) at receptors where CS extract produced > 50% inhibition of radioligand binding. CS extract constituents exhibited >50% binding to a wide range of addiction-related receptors, including monoamine, opioid, benzodiazepine, glucocorticoid, cannabinoid, and orexin receptors, among others. In contrast, nicotine alone and EA extract did not produce > 50% binding at any non-nAChRs, except for alpha 2B receptors and muscarinic M1, M4, and M5 receptor subtypes. Nicotine alone also bound to 5-HT2A receptors and the serotonin transporter. mong the functional analyses, CS extract showed significant enzyme-inhibitory activity at COX-1 and -2 (Figure 7), with CS extract inhibiting COX-1 by 74% and COX-2 by 107% at 1 µM nicotine (EC50 = 0.63 and 0.34 µM, respectively). Nicotine alone showed no activity at either COX-1 or -2. CS extract had much weaker functional effects at other non-cholinergic targets, including agonist effects at adrenoreceptors 1A and 2A (EC50 = 7.9 µM and 3.4 µM, respectively) and antagonist effects at 5HT-2B receptors (EC50 = 7.7 µM). No significant activity of CS extract or nicotine alone was observed at other receptors.

**Figure 7.**
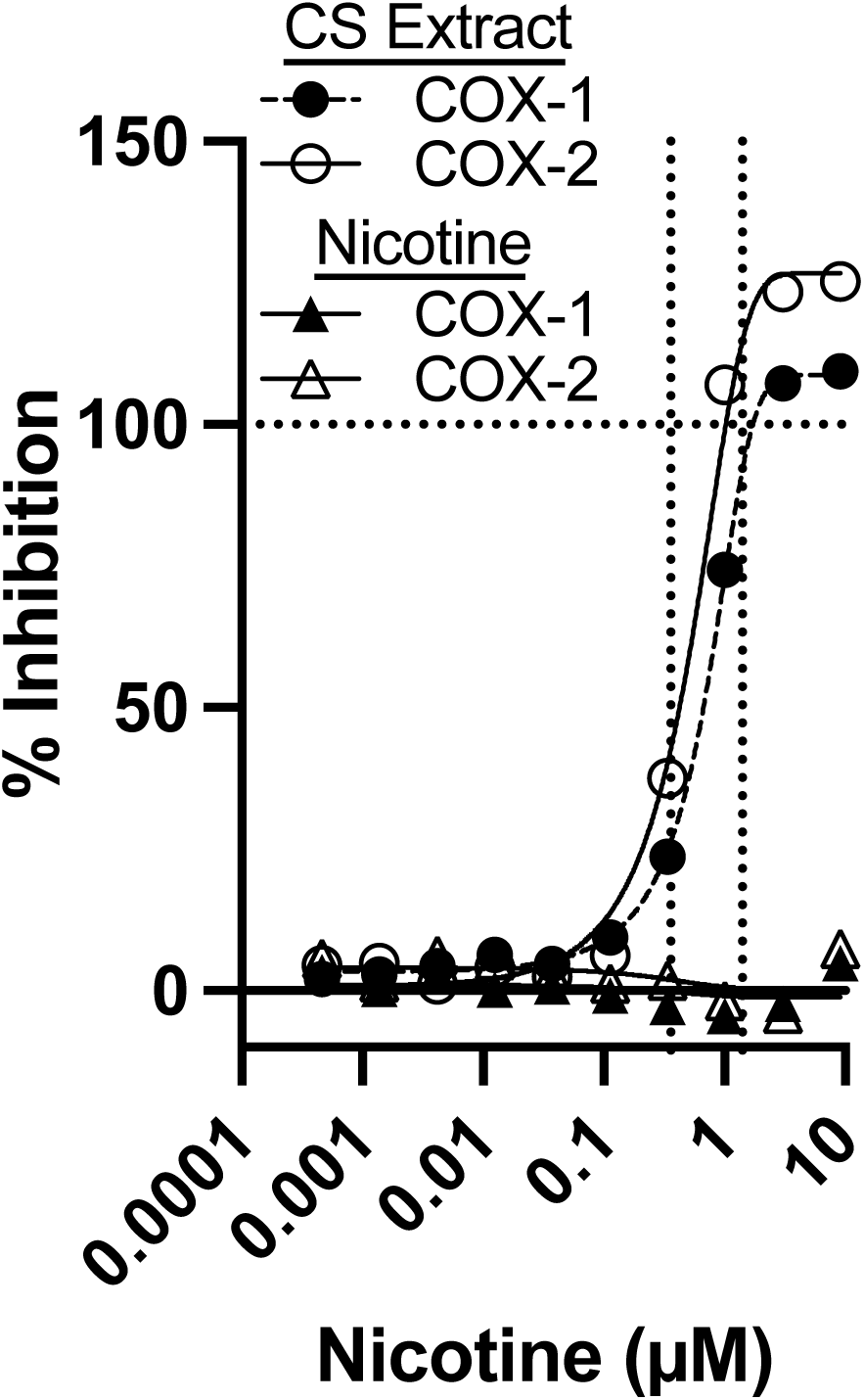
Percent inhibition of COX-1 or -2 enzymes in vitro by CS extract or nicotine alone across a range of nicotine concentrations. Horizontal dotted line indicates 100% inhibition. Vertical dotted lines indicate the estimated range of nicotine concentrations in brain of smokers.

**Table 4.** Percent inhibition in receptor binding assays by nicotine alone and each extract at a high nicotine concentration (184.9 µM, equivalent to 30 µg/kg) at AChRs and non-nAChRs that CS extract produced greater than 50% inhibition. See text for more details and Supplementary Table 1 for a list of all targets tested. Note: ND = not determined.

|  | Percent Inhibition |  |  |
|---|---|---|---|
| **CNS Receptor** | **Nicotine** | **CS Extract** | **EA Extract** |
| Dopamine D1 | 8.99364 | 82.3104 | -2.45 |
| Dopamine D2 | 8.95988 | 84.515 | -11.1 |
| Dopamine transporter | 13.5147 | 70.8817 | 4.27183 |
| Serotonin 5-HT<sub>1A</sub> | 21.582 | 81.4248 | 1.71818 |
| Serotonin 5-HT<sub>2A</sub> | 59.7195 | 82.1143 | -9.77 |
| Serotonin 5-HT<sub>2B</sub> | ND | 90.2139 | 14.0191 |
| Serotonin 5-HT<sub>2C</sub> | 19.5643 | 88.2526 | 29.4568 |
| Serotonin 5-HT<sub>3</sub> | 44.6184 | 64.4183 | 6.22882 |
| Serotonin transporter | 50.6291 | 57.1144 | 34.3662 |
| Adrenergic Receptor (AR) *α*<sub>1A</sub> | 12.311 | 91.0346 | -1.9 |
| AR *α*<sub>2A</sub> | 36.2874 | 91.7365 | 32.6347 |
| AR *α*<sub>2B</sub> | 52.0032 | 88.338 | 50.8925 |
| AR *α*<sub>2C</sub> | 23.9341 | 94.9935 | 39.8256 |
| AR β<sub>2</sub> | 4.29825 | 51.2281 | -2.81 |
| NET transporter | 34.2769 | 96.1373 | 38.9352 |
| Delta opioid | -6.04 | 96.1064 | -25.3 |
| Kappa opioid | 36.7697 | 95.5184 | ND |
| Mu opioid | 26.1211 | 95.1533 | 37.9888 |
| Androgen receptor | -21.7 | 71.9586 | -27.2 |
| Benzodiazepine | -3.72 | 81.019 | -19.6 |
| Cannabinoid CB<sub>2</sub> | 8.0683 | 53.3037 | 15.6436 |
| Cl- channel | -11.4 | 62.0914 | -7.82 |
| Glycine | 18.4582 | 52.9663 | 18.3251 |
| Gluccocorticoid | 13.1608 | 77.7655 | 10.7622 |
| Kainate | 32.5636 | 51.6234 | 38.2535 |
| Na+ channel | -5.01 | 79.9739 | 9.09882 |
| Orexin OX1 | 4.84879 | 74.7816 | -6.35 |
| nAChR *α*4β2 | 100.18 | 99.9205 | 100.291 |
| nAChR *α*7 | 97.0866 | 97.5246 | 87.9946 |
| Muscarine M1 | 55.24 | 97.144 | 55.0761 |
| Muscarine M2 | 43.039 | 97.9279 | 47.3805 |
| Muscarine M3 | 38.3751 | 77.1804 | ND |
| Muscarine M4 | 62.9712 | 84.3681 | 63.8581 |
| Muscarine M5 | 57.7749 | 76.6804 | 56.6462 |

## 4. Discussion

### 4.1. Overall findings

The present study identified several key differences in the behavioral effects of CS extract and nicotine alone, while fewer differences were observed between EA extract and nicotine alone. First, baseline SA was lower and the descending limb of the unit dose-response curve was shifted downward/leftward for CS and EA extracts compared to nicotine alone. Second, compensatory increases in SA during dose reduction were greater and occurred over a wider range of unit doses for CS extract, but not EA extract, than nicotine alone. Third, the reinforcement threshold was lower for CS and EA extract versus nicotine alone in males, while it was only lower for CS extract in females. Fourth, although there were no differences in elasticity of demand between formulations, intensity of demand (Q0) was significantly lower for CS extract, but not EA extract, versus nicotine alone. Fifth, stress-induced reinstatement for CS extract was comparable to nicotine alone, whereas it was weaker for EA extract compared to nicotine alone. Unique neuropharmacological effects of CS extract were also observed. Agonist potency at α4ß2 nAChRs was greater for CS extract than nicotine alone, and CS extract, but not EA extract, showed greater binding than nicotine alone at numerous non-nAChRs relevant to TUD and other substance use disorders. Finally, the present study is the first to show that CS extract, but not nicotine alone, inhibits COX-1 and -2 enzymes at smoking-relevant concentrations in vitro. Together, these findings demonstrate that non-nicotine constituents in CS may collectively influence the reinforcing effects of combusted cigarettes, while they may have relatively less influence on the reinforcing effects of ENDS. The present findings are consistent with prior studies showing greater reinforcing and other behavioral effects of CS and EA extracts versus nicotine alone and provide new insight into potential mechanisms mediating these effects.

### 4.2. Behavioral mechanisms of non-nicotine constituent effects

The present findings indicate that non-nicotine constituents primarily increased the reinforcing <u>potency</u> of CS extract, as indicated by lower baseline SA, a downward/leftward shift in the descending limb of the dose-response curve, and a lower reinforcement threshold in each sex for CS extract compared to nicotine alone. From this perspective, satiety is achieved with fewer infusions of extract than nicotine alone. In contrast, the reinforcing <u>efficacy</u> of CS extract did not differ significantly from nicotine alone in terms of elasticity of demand. These findings are consistent with other studies that showed greater potency of CS extract under small fixed-ratio schedules, but no difference in reinforcing efficacy (i.e., breaking-point) under progressive-ratio schedules (Brennan et al., 2015; Costello et al., 2014; Marusich et al., 2019). However, the greater potency of CS extract in other studies was indicated by <u>higher</u> rates of CS extract self-administration at unit doses below the training dose, which is the opposite of what we observed. The reason for this difference is unclear, but might be due to several factors. First, we used a different tobacco product and extraction method that likely yielded a chemical profile that differed significantly from extracts used by others, and some studies have shown that CS extract effects can depend on the tobacco product used (Brennan et al., 2015; Marusich et al., 2019). Given that our extraction method was designed to include gas-phase and non-aqueous compounds, levels of certain addiction-related constituents (e.g. acetaldehyde, acetone, etc.) were likely higher than in the aqueous extracts used by others. For example, acetaldehyde levels were 16.9 µg/kg/infusion in the present study, compared to approximately 3 µg/kg/inf in another study that used the same nicotine SA training dose (Marusich et al., 2019). Second, the training dose and/or infusion parameters, such as infusion speed and timeout interval, in the present study differed from some other studies. These parameters can influence the shape and position of the nicotine SA dose-response curve and the underlying neural mechanisms involved (Sorge and Clarke, 2009). Third, the dose-response curve for nicotine alone in some prior studies was unusually flat, indicating an insensitivity to nicotine dose. In addition, unit doses of nicotine alone that normally maintain SA failed to do so in some studies. Fourth, the experimental design of other studies differed from the present study (e.g., using nicotine self-administration or food training before extract exposure, use of different response topographies (lever press vs nose poke).

Although the primary measure of reinforcing efficacy, elasticity of demand, did not differ between formulations, differences in other measures suggest somewhat greater reinforcing efficacy of CS extract compared to nicotine alone. For example, compensatory increases in SA during dose reduction (i.e. increases in unit price) were greater for CS extract than nicotine alone (Figure 2B). Because such compensation reflects an increase in effort expended to defend baseline nicotine intake as unit price increases, compensation could be considered another behavioral measure of reinforcing efficacy, akin to Omax (Hursh, 1991; Hursh and Silberberg, 2008) or breaking point (Stafford et al., 1998). Although Omax did not differ between formulations, this measure refers to a single point on the response output curve at which demand becomes relatively elastic (i.e., Pmax). In contrast, the main effect of formulation on compensation in the present study includes all unit prices/doses, allowing comparison of the overall degree of increase in response output for each formulation. In addition, the higher Pmax for CS extract was nearly statistically significant (p = 0.05). Taken together, these data suggest that the reinforcing efficacy of CS extract was somewhat greater than nicotine alone.

Greater aversive/punishing effects may also mediate the lower responding for higher unit doses of extracts, as the same dose of nicotine can have both reinforcing and punishing effects (LeSage, 2022). We have shown that CS extract, but not EA extract, can produce greater aversive effects than nicotine alone in intracranial self-stimulation and conditioned place aversion models (Harris et al., 2022). However, this was observed at a high nicotine dose (≥ 1 mg/kg, s.c.) that produces serum nicotine concentrations much higher than what is achieved via self-administration of a 30 µg/kg nicotine unit dose under limited access conditions (Craig et al., 2014; LeSage et al., 2002). Nonetheless, nicotine can produce aversive effects at lower doses in other models (e.g. conditioned taste or place aversion induced by noncontingent or self-administered nicotine (Han et al., 2017; C. Liu et al., 2022; Pescatore et al., 2005; Rinker et al., 2008)). However, other studies showing higher or no difference in PR breakpoints for CS extracts compared to nicotine alone, and the greater compensation and trend toward a higher Pmax in the present study, suggests CS extract had greater reinforcing rather aversive/punishing potency at unit doses on the descending limb of the dose-response curve.

### 4.3. Potential neuropharmacological mechanisms of non-nicotine constituent effects

Activation of α4β2 nAChRs plays a critical role in nicotine reinforcement and TUD. The greater potency of CS extract than nicotine alone in activating α4β2 nAChRs is consistent with the greater reinforcing potency of CS extract than nicotine alone in the present study. It may also be relevant to our prior finding that a high dose of the same CS extract produced greater elevations in ICSS thresholds than nicotine alone, indicating greater aversive/anhedonic effects (Harris et al., 2022). Plausible contributors include minor tobacco alkaloids present in CS extract, such as nornicotine, anabasine, and anatabine, which can activate α4β2 nAChRs (Harris et al., 2015).

Although levels of these alkaloids are substantially lower than nicotine in the extract and generally exhibit lower potency and/or efficacy at α4β2 nAChRs, their combined activity— potentially together with that of other known or unknown CS constituents—could have contributed to the greater α4β2 potency of the extract. Greater reinforcing effects of cocktails of nicotine and minor alkaloids compared to nicotine alone observed in some studies support this notion.(Clemens et al., 2009) Further testing this possibility should include future studies using bioassay-guided fractionation to identify active extract fractions (e.g., Noya et al., 2013) and/or comparing nicotine alone to nicotine combined with one or more other constituents identified in those fractions.

The present study is the first to examine binding and functional effects of extracts at a wide range addiction-related noncholinergic receptors. At the high concentration used for initial screening, CS extract, but not EA extract or nicotine alone, produced ≥50% inhibition of ligand binding at numerous non-nAChR targets, including dopamine, serotonin, adrenergic, and opioid receptors and transporters. However, the nicotine-equivalent concentration used in this screen substantially exceeded estimated brain nicotine concentrations in smokers and in rats self-administering the training dose used here. Although this in vitro to in vivo comparison can’t be made directly without accounting for pharmacokinetic factors (e.g., protein binding and tissue distribution, Yoon et al., 2012), such a large difference suggests the screening results are likely not representative of in vivo target engagement. Accordingly, the follow-up functional assays identified activity at only a subset of targets, with COX-1 and COX-2 inhibition being the most notable non-nAChR effects observed at nicotine-equivalent concentrations closer to those expected in vivo. Thus, the binding findings simply illustrate the diversity of pharmacological targets that non-nicotine constituents in CS can engage rather than evidence that these constituents could be involved in typical smoking or CS extract self-administration. However, they may be useful for identifying CS constituents with a specific mechanism of action to study further.

CS extract, but not nicotine alone, inhibited COX-1 and COX-2 at nicotine concentrations corresponding to smoking-relevant nicotine levels. This extends a prior report that nicotine-and a tar-free gas-phase CS extract suppressed platelet thromboxane production through COX-1 inhibition and directly inhibited both COX isoforms in vitro (Kashiwagi et al., 2017). COX-1 and COX-2 contribute to neuroinflammatory signaling by converting arachidonic acid to PGH₂, the precursor of prostanoids including PGE₂, which can regulate microglial activation and the production of inflammatory cytokines and chemokines (Fitzpatrick, 2004). Neuroinflammation may be relevant to TUD because human imaging studies have found lower binding to a putative marker of neuroimmune activation in smokers (Brody et al., 2025a, 2025b, 2017), whereas nicotine withdrawal increases neuroinflammatory signaling in preclinical models (Soares and Picciotto, 2024). Withdrawal-associated neuroinflammation may, in turn, contribute to affective and cognitive symptoms that promote craving and relapse (Kohno et al., 2019; Soares and Picciotto, 2024). Although nicotine’s activation of α7 nAChRs has often been invoked to explain anti-inflammatory effects associated with tobacco exposure,(Soares and Picciotto, 2024) evidence suggests that non-nicotine constituents may also contribute (Brody et al., 2018, 2017). The present finding identifies COX inhibition as another potential non-nicotine mechanism linking cigarette smoke exposure to neuroimmune modulation. Future studies identifying the responsible constituents and directly manipulating COX signaling are needed to determine whether COX inhibition contributes to the behavioral or neuroimmune effects of CS extract.

Our in vivo microdialysis findings suggest that differential NAc dopamine release did not contribute to the differences in behavioral effects of CS and EA extracts versus nicotine alone at an acute level. Responses to both extracts were nevertheless more variable overall than responses to nicotine alone, particularly at later time points. This pattern could reflect heterogeneous dopaminergic responses to non-nicotine constituents, although other sources of variability cannot be excluded. A limitation of our approach is that the 20-min collection intervals may have obscured brief or temporally shifted effects. Approaches with greater temporal resolution, such as fast-scan cyclic voltammetry, may be better suited to detecting differences in rapid dopamine dynamics. Measuring NAc dopamine release during response-contingent self-administration would also more closely model the conditions under which behavioral differences between formulations were observed. This latter point is particularly important, as the current experimental paradigm for microdialysis was based on the delivery of two acute i.v. drug infusions. Thus, future studies are needed to assess the impacts of repeated drug exposures within and across days, which would better mimic drug self-administration paradigms.

### 4.4. Reinstatement effects

Reinstatement for all three formulations was induced by yohimbine, but was weaker for EA extract compared to nicotine alone. The similarity of yohimbine effects between CS extract and nicotine alone may be due to extinction and reinstatement tests occurring in the <u>presence</u> of infusion-associated cues, as a prior study showed greater yohimbine reinstatement of CS extract responding than nicotine alone only when extinction and reinstatement tests occurred in the <u>absence</u> of cues (Costello et al., 2014). However, the effects of yohimbine, cues, and other reinstatement stimuli (e.g. nicotine or CS extract priming) have been inconsistent across studies (Costello et al., 2014; Cross et al., 2020; Gellner et al., 2023). Regardless, the present findings suggest that stress-induced reinstatement may not be affected by a history of exposure to non-nicotine constituents in CS. To our knowledge, the present study is the first to examine reinstatement of responding in rats trained for EA extract SA. The weaker effect of yohimbine on EA extract responding compared to nicotine alone suggests that stress-induced reinstatement of EA extract responding may be attenuated by exposure to non-nicotine constituents in EA extract. Studies using other stressors, such as footshock or social defeat, are needed to determine whether the attenuation observed following EA extract self-administration generalizes beyond yohimbine.

### 4.5. Sex differences

To our knowledge, the present study is the first to examine sex differences in the reinforcing effects of CS and EA extracts. Baseline intake of nicotine alone was higher in females than males, consistent with prior reports (Donny et al., 2000; Grebenstein et al., 2013; Klein et al., 2004; Lynch, 2009; Park et al., 2007). This may be due to sex differences in nicotine pharmacokinetics, such as faster nicotine clearance or higher peak brain concentrations in females (Klein et al., 2004; LeSage et al., 2007; Rosecrans, 1972). There was a lower threshold for both extracts compared to nicotine alone in males, but only for CS extract in females. Males, but not females, also showed greater compensation for CS extract than nicotine alone over a wider range of unit doses compared to nicotine alone. However, the magnitude of compensation for nicotine alone was similar between sexes, which contrasts with our prior study showing less compensation in females than males.(Grebenstein et al., 2013) This may be due to the shorter session duration in the present (2 hr) compared to the prior study (23 hr). Overall, the findings raise the possibility of greater sensitivity of males to non-nicotine constituents. However, this study was not powered specifically to detect sex differences because sex effects were not a primary interest. Future studies are needed to examine the reliability of these observed sex effects.

### 4.6. Limitations

The lack of leftward shift in the ascending limb (1 to 4 µg/kg) of the dose-response curve is consistent with the prior studies showing no difference in responding for CS extract and nicotine alone in this dose range (Costello et al., 2014; Marusich et al., 2019). This is likely due to the reduction in concentration of non-nicotine constituents (and their behavioral effects) as the extract is diluted to reach the desired nicotine concentration. As such, the concentration of non-nicotine constituents at the higher nicotine unit doses (7.5 to 30 µg/kg), which can produce serum nicotine concentrations in the range of smokers/vapers, are more relevant to non-nicotine constituent exposure in humans. The inability to manipulate the nicotine concentration independently of non-nicotine constituents is a major limitation of the present and all prior behavioral studies of tobacco product extracts. Holding the concentration of non-nicotine constituents relatively constant as the nicotine unit dose is reduced would better clarify how non-nicotine constituents can influence the reinforcing potency of nicotine, because their effects would be more likely to manifest at low nicotine doses. As such, the nicotine reinforcement threshold CS extract in that context may be lower than observed in the present study. This could be achieved using CS extracts from very low nicotine content (VLNC) cigarettes(Ding et al., 2017; Richter et al., 2016) and “spiking” the extracts with pure nicotine to achieve the desired nicotine unit doses. This would have important FDA tobacco regulatory implications, as it would provide a more valid preclinical model of smoking VLNC cigarettes and might support setting a low nicotine standard for cigarettes than self-administration models that use nicotine alone.(DHHS and FDA, 2018; LeSage et al., 2018)

### 4.7. Conclusions

Despite its limitations, the present study is a significant advance in preclinical research on the addiction-related effects of tobacco product extracts. It provides the most extensive dose-response analysis to date, allowing comparison of reinforcement thresholds and elasticity of demand to parse effects of non-nicotine constituents on the reinforcing potency versus efficacy of CS extract, respectively. Although it may be underestimated, our finding of a lower reinforcement threshold for CS extract compared to nicotine alone has important implications for preclinical tobacco regulatory science and associated FDA tobacco control policy. Future studies are needed to examine whether the greater α4β2 agonist potency, MAO inhibition, and COX inhibition observed *in vitro* with CS extract in this or our previous study (Harris et al., 2022) occur *in vivo* and mediate the behavioral effects of CS extract.

## Acknowledgements

The authors thank Megan McGuigan and Dr. Clifford Watson at the Tobacco and Volatiles Branch, Division of Laboratory Sciences, National Center for Environmental Health, Center for Disease Control (Atlanta, GA) for measuring levels of toluene, benzene, and ethylbenzene in extract samples. We also thank Dr. Fred Apple and Julie Kloss from Hennepin Healthcare (Minneapolis, MN) for conducting the glycol analyses.

