## Supplementary Table and Figures for "The collective effect of non-nicotine constituents on the reinforcing and neuropharmacological effects of comprehensive cigarette smoke and electronic nicotine delivery system (ENDS) aerosol extracts in rats"

**Supplementary Table 1.** Receptor/channel/enzyme targets for functional assays.

|  | **Control Compound** | |  |
| --- | --- | --- | --- |
| **Receptor/Channel/Enzyme** | **Agonist/Opener** | **Antagonist/Inhibitor** | **Assay Type** |
| Acetylcholine ⍺4ß2 | Nicotine | Dihydro-ß-erythroidine | Ion Channel |
| Acetylcholine M1 | Acetylcholine | Atropine | Calcium Flux |
| Acetylcholine M2 | Acetylcholine | Atropine | Calcium Flux |
| Acetylcholine M3 | Acetylcholine | Atropine | Calcium Flux |
| Acetylcholinesterase | | Physostigmine | Enzymatic |
| Adenosine A2A | NECA | SCH 442416 | Calcium Flux |
| Adrenergic A1A | A61603 Hydrobromide | Tamsulosin | Calcium Flux |
| Adrenergic A2A | UK 14304 | Yohimbine | cAMP |
| Adrenergic B1 | (1)-Isoproterenol | Betaxolol | cAMP |
| Adrenergic B2 | (1)-Isoproterenol | ICI 118,551 HCL | cAMP |
| Androgen | BMS-564929 | Geldanamycin | NHR Nuclear Translocation |
| Calcium Channel V1.2 | | Isradipine | Ion Channel |
| Cannabinoid 1 | CP 55940 | AM 251 | cAMP |
| Cannabinoid 2 | CP 55940 | SR 144528 | cAMP |
| Cholecystokinin A | Sincalide | SR 27897 | Calcium Flux |
| Cyclooxygenase (COX-1) | | Indomethacin | Enzymatic |
| Cyclooxygenase (COX-2) | | NS-398 | Enzymatic |
| Dopamine D1 | Dompamine | SCH 39166 | cAMP |
| Dopamine D2 | Dopamine | Resperidone | cAMP |
| Dopamine Transporter | | GBR-12909 | Transporter |
| Endothelin A | Enthothelin 1 | BMS 182874 | Calcium Flux |
| GABA A | GABA | Picrotoxin | Ion Channel |
| Glucocorticoid | Dexamethasone | Mifepristone | NHR Protein Interaction |
| hERG |  | Astemizole | Ion Channel |
| Histamine 1 | Histamine | Mepyramine | Calcium Flux |
| Histamine 2 | Histamine | Tiotidine | cAMP |
| Insulin |  | BMS-754807 | Kinases |
| K Channel KvLQT1/minK | ML-277 | XE 991 | Ion Channel |
| Monamine Oxidase (MAOA) | | Clorgyline | Enzymatic |
| NAV1.5 |  | Lidocaine HCL | Ion Channel |
| NMDA 1A/2B | L-Glutamic Acid | (+)-MK 801 | Ion Channel |
| Norepinephrine Transporter | | Desimpramine | Transporter |
| Opioid D1 | DADLE | Naltrinben | cAMP |
| Opioid K1 | Dynorphin A (1-17) | nor-Binaltorphimine | cAMP |
| Opioid M1 | DAMGO | Naloxone | cAMP |
| Phosphodiesterase 3A | | Cilostamide | Enzymatic |
| Phosphodiesterase 4D2 | | Cilomilast | Enzymatic |
| Rho kinase (ROCK1) | | Staurosporine | Kinases |
| Serotonin 1A | Serotonin | Spiperone | cAMP |
| Serotonin 1B | Serotonin | SB 224289 | cAMP |
| Serotonin 2A | Serotonin | Altanserin | Calcium Flux |
| Serotonin 2B | Serotonin | LY 272015 | Calcium Flux |
| Serotonin 3A | Serotonin | Bremestron | Ion Channel |
| Serotonin Transporter | | Clomipramine | Transporter |
| Tyrosine kinase (LCK) | | Gleevec | Kinases |
| Vasopressin 1A | [Arg8]-Vasopressin | SR 49059 | Calcium Flux |
| VEGFR2 |  | SU-11248 | Kinases |

**Supplementary Figure 1.** Mean (±SEM) number of infusions in rats responding for CS extract, EA extract, nicotine alone, or saline during the first ten sessions of acquisition of SA at FR 1 and the final three sessions of stability at each FR.

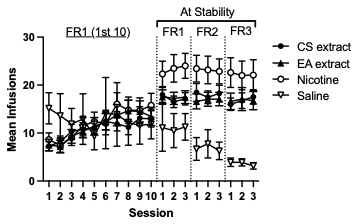

**Supplementary Figure 2.** Mean (±SEM) number of responses on the active and inactive levers during the course of unit dose reduction in rats responding for nicotine alone (left panel), CS extract (center panel), or EA extract (right panel).

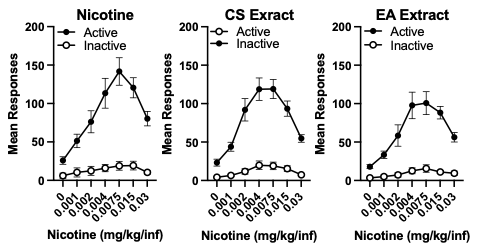

**Supplementary Figure 3.** Mean (±SEM) number of infusions expressed as a percentage of baseline (B, 30 µg/kg unit dose) during the first 10 sessions of extinction and the final three sessions when extinction criteria were met (F) .

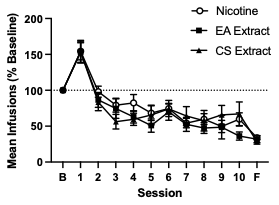

**Supplementary Figure 4.** Mean (±SEM) number of infusions in rats responding for saline + ETOH that met acquisition criteria. Each point represent the number of infusions at each ETOH concentration that matched the ETOH concentrations in each unit dose of nicotine alone, CS extract, or EA extract.

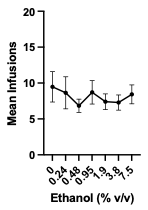
